# Mechanistic basis of staphylococcal interspecies competition for skin colonization

**DOI:** 10.1101/2023.01.26.525635

**Authors:** Joseph J. Maciag, Constance Chantraine, Krista B. Mills, Rahul Yadav, Alexander E. Yarawsky, Catherine T. Chaton, Divya Vinod, Nicholas C. Fitzkee, Marion Mathelié-Guinlet, Yves F. Dufrêne, Paul D. Fey, Alexander R. Horswill, Andrew B. Herr

## Abstract

Staphylococci, whether beneficial commensals or pathogens, often colonize human skin, potentially leading to competition for the same niche. In this multidisciplinary study we investigate the structure, binding specificity, and mechanism of adhesion of the Aap lectin domain required for *Staphylococcus epidermidis* skin colonization and compare its characteristics to the lectin domain from the orthologous *Staphylococcus aureus* adhesin SasG. The Aap structure reveals a legume lectin-like fold with atypical architecture, showing specificity for N-acetyllactosamine and sialyllactosamine. Bacterial adhesion assays using human corneocytes confirmed the biological relevance of these Aap-glycan interactions. Single-cell force spectroscopy experiments measured individual binding events between Aap and corneocytes, revealing an extraordinarily tight adhesion force of nearly 900 nN and a high density of receptors at the corneocyte surface. The SasG lectin domain shares similar structural features, glycan specificity, and corneocyte adhesion behavior. We observe cross-inhibition of Aap- and SasG-mediated staphylococcal adhesion to corneocytes. Together, these data provide insights into staphylococcal interspecies competition for skin colonization and suggest potential avenues for inhibition of *S. aureus* colonization.

---

The skin microbiome plays an important role in the maintenance of homeostasis and contributes to localized immune responses. Commensal skin bacteria such as *Staphylococcus epidermidis* generally play a beneficial role; for example, *S. epidermidis* and other coagulase-negative staphylococci can suppress invasion by *S. aureus* or other pathogens directly, via secreted factors ^1–5^, or indirectly, by cooperating with local immune cells in the skin to induce antimicrobial protein production by keratinocytes ^6^. *S. epidermidis* is a near-ubiquitous colonizer of healthy skin, whereas *Staphylococcus aureus* typically colonizes the skin in ~5% of healthy individuals ^7–10^. However, up to 90% of individuals with atopic dermatitis (AD) are colonized with *S. aureus*, and AD flares are characterized by dysbiosis of the skin microbiome and overgrowth of *S. aureus* to a striking degree ^11^. *S. aureus* colonization of the skin is associated with decreased epidermal barrier function and increased severity of inflammatory outcomes in AD ^12–17^ and AD patients often progress to other, more threatening, atopic diseases including food allergy, allergic rhinitis, and asthma ^13, 14, 18^.

Cell wall surface proteins of Gram-positive bacteria mediate a wide variety of functions related to colonization and virulence. In both *S. epidermidis* and *S. aureus*, a large number of cell wall-anchored (CWA) proteins containing a C-terminal LPXTG sequence motif are covalently attached to peptidoglycan in the cell wall; these CWA proteins mediate processes including adhesion to host tissue, immune evasion, biofilm formation, and a variety of other enzymatic and non-enzymatic functions ^19–23^. *S. epidermidis* expresses a CWA protein known as the Accumulation-associated protein (Aap), a large, multidomain protein first identified by its role in the accumulation phase of biofilm formation ^24^. In addition to its role in mediating intercellular adhesion in biofilms ^25–31^, Aap has been shown to be essential for skin colonization by *S. epidermidis* via adhesion to healthy human corneocytes ^32–34^.

The domain architecture of Aap includes the N-terminal A domain, comprising the disordered A-repeat region with 11 imperfect repeats of a 16-residue motif and the following lectin domain. The A domain is immediately followed by 5 to 17 large 128-residue B-repeats that adopt an unusual, highly extended fold made up of unsupported 3-stranded β-sheets with minimal hydrophobic side chain packing ^26, 35^. Each B-repeat is comprised of a G5 subdomain followed by a smaller E (or spacer) subdomain with a similar fold ^26, 35, 36^. The B-repeat superdomain terminates with a variant G5 domain. The C-terminal end of Aap contains a repeating proline/glycine-rich (PGR) sequence that functions as a highly extended, noncompressible stalk region ^37^. The LPXTG anchor is found at the end of the PGR region and is covalently attached to the cell wall by Sortase A ^38–40^. The B-repeat superdomain is the region responsible for the intercellular adhesion leading to biofilm formation, but it first requires proteolytic processing by the trypsin-like staphylococcal protease SepA to remove the A domain (which includes the lectin domain) ^41^. Once processed, the B-repeat superdomain can engage in self-assembly events mediated by free Zn^2+^ to initially form antiparallel dimers that create twisted rope-like strands between staphylococcal cells in the nascent biofilm ^25, 26, 35, 42–44^. These dimeric structures can further assemble to form tetrameric species that then nucleate functional amyloid fibers within the biofilm ^27, 43, 45^.

While the B-repeat superdomain mediates intercellular adhesion in the biofilm, the A domain plays an important role in adhesion to both biotic and abiotic surfaces ^32–34^. Early work demonstrated that the A domain was necessary for adhesion to human corneocytes ^33^, and it has also been implicated in biofilm attachment to a catheter surface in a rat infection model ^28^. Recently, it was demonstrated by Roy et al. that the Aap lectin subdomain within the A domain is the critical component required for skin colonization, as it is necessary and sufficient for *S. epidermidis* adhesion to healthy human corneocytes ^34^. The authors also demonstrated that *S. epidermidis* adhesion could be blocked by enzymatic cleavage of specific glycan moieties, confirming that lectin binding activity was necessary for adhesion ^34^.

*S. aureus* expresses an Aap ortholog named SasG that adopts a near-identical domain architecture including A-repeats, lectin domain, B-repeat superdomain, Pro/Gly-rich stalk, and LPXTG sortase motif (Fig. S1) ^46^. Similar to Aap, the SasG A domain mediates adhesion to desquamated nasal epithelial cells ^47, 48^. Likewise, the B-repeat superdomain of SasG adopts the same fold as Aap B-repeats and can self-assemble in a Zn^2+^-dependent manner to mediate both homophilic and heterophilic interactions (with Aap B-repeats) ^36, 44, 49^. Many unresolved questions remain about the structure of the Aap and SasG lectin domains, the relative conformation of the lectin with respect to the subsequent B-repeat superdomain, the native ligand specificity of both lectins, and the potential for competition between the two staphylococcal species for colonization of the human skin niche. Here we present multidisciplinary data describing high-resolution structures of both Aap and SasG lectin domains, the dynamics of the lectin fold, specificity for glycan ligands, adhesion to healthy human skin corneocytes, force spectroscopy on the adhesive interaction, and evidence for direct competition between *S. aureus* and *S. epidermidis* for adhesion to corneocytes and colonization of skin.

## Results

### Aap lectin adopts a legume lectin-like fold with an atypical binding site architecture

The crystal structure of Aap lectin was solved to 1.06 Å resolution; phase determination involved single-wavelength native anomalous diffraction from the δ sulfur atoms of two methionine residues (M436 and M593) and a structural calcium ion. Aap lectin adopts a fold similar to a large number of legume lectins, with two 6- or 7-stranded antiparallel β-sheets bridged by an orthogonal 5-stranded sheet along with several long loops, some of which include short α-helices and one 3_10_ helix (Fig. 1a-b). The 6-stranded β-sheet is relatively flat and solvent-exposed, transitioning on one end into the orthogonal 5-stranded sheet, whereas the 7-stranded sheet forms a concave surface over which four loops curl inward, coming together to create the boundaries of a large pocket on the protein surface with a tyrosine (Y580) at its base (Fig. 1d). Two structural metal ions, a Ca^2+^ and a Na^+^, are bound on either side of the Y580 pocket. The Aap fold is most closely related to the lectin domain of SraP, another staphylococcal surface protein ^50^; like SraP, Aap is a monomeric lectin (Fig. S2) in contrast to most of the legume lectins that form dimers or higher-order multimers. The location of the putative Aap binding pocket anchored by Y580 is located centrally with regard to the concave β-sheet, which differs from the more distal glycan binding site of SraP and other legume lectins (Fig. 1d-e).

**Fig. 1:**
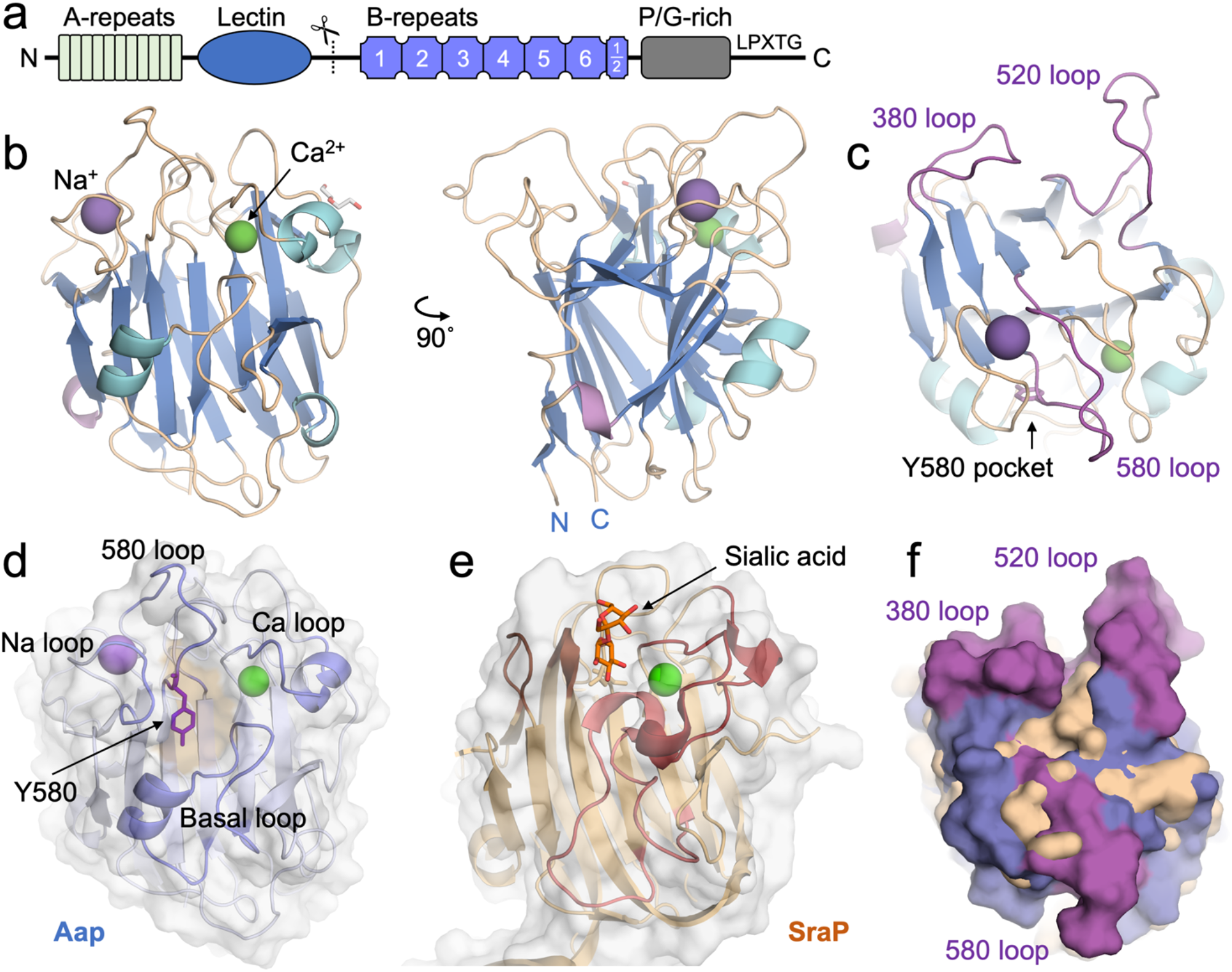
Crystal structure of Aap lectin reveals an atypical legume lectin-like fold. (**a**) Domain arrangement of Aap from *S. epidermidis* 1457. The scissors icon indicates the cleavage site for SepA that unmasks the B-repeat superdomain and allows protein-dependent biofilm formation. (**b**) Front and side views of the crystal structure of Aap lectin domain, colored by secondary structural elements. β-strands are colored blue, α-helices cyan, 3_10_ helices lavender, and loops tan. (**c**) Top view of Aap lectin, highlighting the divergent long loops (380, 520, and 580 loops) in purple. (**d**) Front view of Aap lectin with transparent surface, showing the loops that converge on the front face to create a putative glycan binding pocket with Y580 forming its floor. The pocket is colored tan to aid in visualization. (**e**) Front view of the related lectin domain from the *S. aureus* adhesin SraP, showing the equivalent loops that form its binding pocket, which is located further toward the distal end of the domain compared to Aap. (**f**) Surface view of the top of the lectin domain of Aap superimposed on SraP. Aap is colored in blue, with the divergent long loops colored purple; SraP is colored tan. The long loops extend beyond the SraP boundary, creating a distinct surface that may be used for target receptor recognition.

Legume lectins are characterized by a structural Ca^2+^ ion adjacent to a conserved central aspartate residue with a *cis* backbone configuration whose side chain is positioned to interact with the glycan ligand ^51, 52^. In the case of Aap, the conserved Ca^2+^ ion is present; coordination of this Ca^2+^ ion constrains one of the loops (‘Ca loop’) that folds over the concave β-sheet and helps create the Y580 pocket (Fig. 1d, Fig. 2a). The conserved aspartate (D431 in Aap) is also present and forms a water-mediated interaction with the neighboring Ca^2+^ ion; however, D431 is in the *trans* configuration in Aap (Fig. 2a-b). In other legume lectins, this conserved aspartate engages the glycan ligand, but D431 in Aap instead forms an H-bond with the backbone nitrogen of Y580, stabilizing another loop (‘580 loop’) that curls in over the concave β-sheet to help define the binding pocket (Fig. 1d, Fig. 2a). Interestingly, on the opposite side from the Ca^2+^ ion, Aap also features a structural Na^+^ ion with a coordination number of 5, ligated by 3 backbone carbonyl oxygens, a water, and the side chain oxygen of Q579. Thus, the coordination shells of the Ca^2+^ and Na^+^ ions provide constraints from both sides that stabilize the unusual configuration of residues Q579-Y580, positioned where the glycan typically binds in other legume lectins (e.g., see transparent sialic acid, Fig. 2b). As a result of these constraints provided by the metal ions, the surface in this region of Aap lectin forms a large pocket with a solvent-accessible volume of 125 Å^3^ that contains the aromatic ring of Y580 centered in the pocket floor. The walls of the pocket are created by a convergence of loops including the Ca^2+^-bound loop, the ‘580 loop’ downstream of the bend at Y580, and part of the Na^+^-bound loop; the bottom portion of the pocket is defined by the ‘basal loop’, a long loop and α-helix ‘plug’ extending from the base of the β-sheet that folds up across the lower face of the lectin. In contrast, although SraP has a similar loop/helix plug defining the lower boundary of the binding site, in SraP it extends further up to create a binding pocket closer to the top of the lectin domain compared to Aap (Fig. 1d-e). Furthermore, the SraP loops corresponding to Aap’s 580 loop and the Na^+^-bound loop are significantly shorter and are not long enough to curl over the concave β-sheet, providing a shallower surface that accommodates the bound glycan ligand (Fig. 1e).

**Fig. 2:**
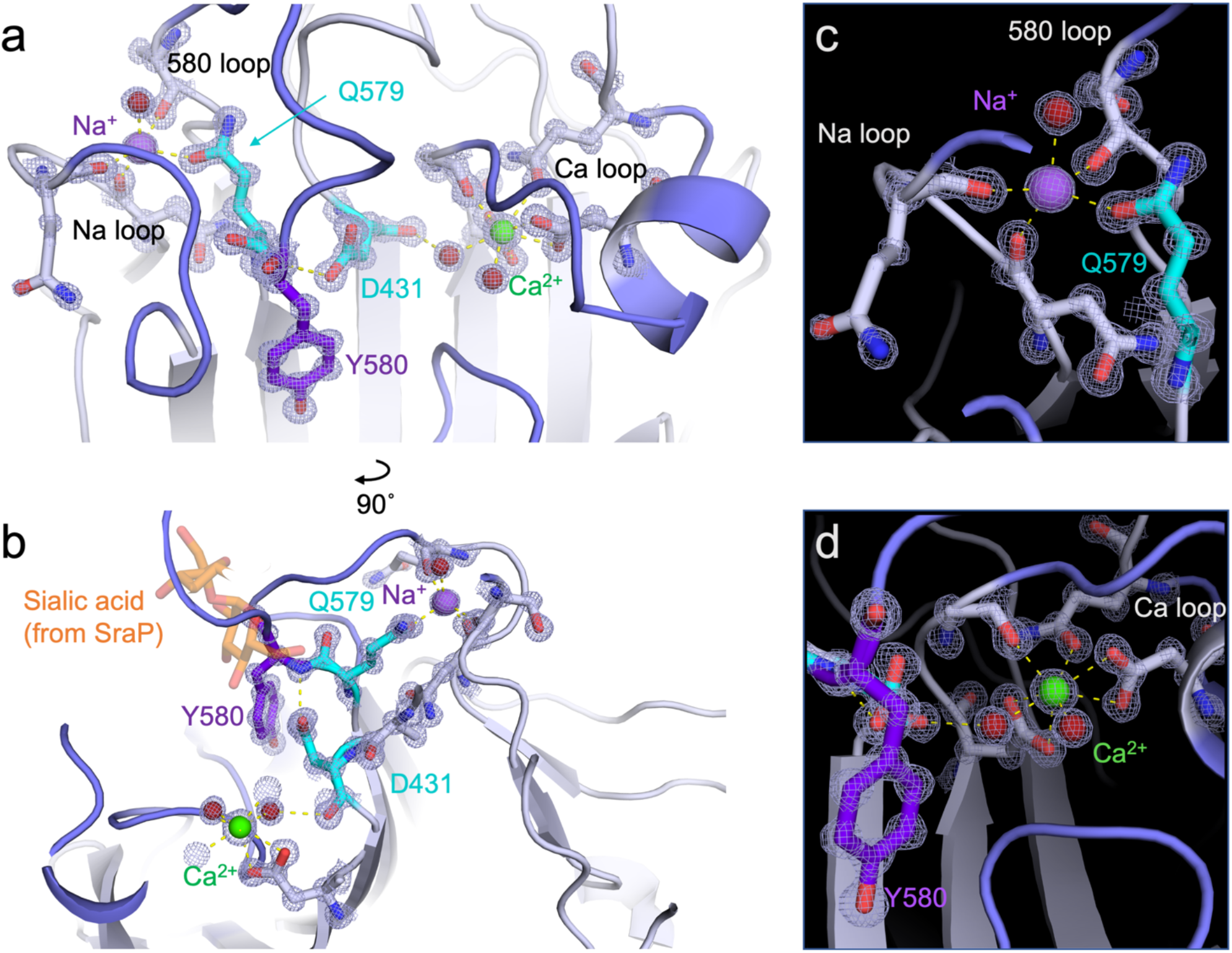
Atypical architecture near the Aap glycan binding site. (**a**) Model and 2fo-fc electron density map illustrating the front view of the Aap lectin in the region of the binding site. The main chain forms a sharp bend at Q579-Y580 that helps create the binding pocket with Y580 as its floor; this bend in the main chain is stabilized on either side via interactions with structural metal ions. D431 is the conserved aspartate residue from legume lectins that typically forms a *cis* peptide bond and whose side chain usually interacts with the glycan ligand. In Aap, D431 forms a *trans* peptide bond and the side chain interacts with the amide nitrogen of Y580, stabilizing the floor of the atypical binding pocket. The carbonyl oxygen of D431 interacts with the structural Ca^2+^ ion indirectly via a water-mediated H-bond. Q579 itself forms a direct H-bond with the bound Na^+^ ion on the opposite side of the domain from the Ca^2+^ binding site. (**b**) View from behind the Ca^2+^ ion looking toward the binding pocket, showing the interaction network involving both Ca^2+^ and Na^+^ ions, D431, and the Q579-Y580 region of the main chain. For reference, the sialic acid from the SraP binding site is shown as transparent sticks, showing that Y580 occupies the same approximate position as bound glycans in other related lectins. (**c**) Details of the Na^+^ coordination scheme, which shows a typical coordination number of 5. (**d**) Details of the Ca^2+^ coordination scheme, with a typical coordination number of 8.

### Solution structure of the lectin-B-repeat junction and lectin domain dynamics

The N- and C-termini of the Aap lectin domain are adjacent to one another (Fig. 1b), which will direct the intrinsically disordered A-repeat region back toward the B-repeat superdomain. As a result, the lectin domain would be found at the extreme terminus of the highly elongated B-repeat superdomain, consistent with its function in adhesion to host tissue. In order to understand the structural arrangement between the lectin and the B-repeat region, we expressed a lectin-Brpt construct containing the native Aap sequence (351-816) of the lectin through the first 1.5 B-repeats (i.e., the 1^st^ two G5 subdomains and the intervening E or spacer subdomain). This B-repeat length corresponds to a stable construct from the C-terminal portion of the B-repeat superdomain that has been characterized extensively ^25–27, 35, 42, 43, 45^. The lectin domain alone, the lectin-Brpt construct, and the corresponding N-terminal Brpt1.5 region alone were characterized in solution by small-angle X-ray scattering (SAXS) (Fig. 3a). A dimensionless Kratky plot was consistent with a globular fold for the lectin domain and elongated conformations for both lectin-Brpt and Brpt1.5 alone (Fig. 3b). The lectin domain alone behaved as a globular protein, as expected, with a D⊓a× of 56 Å from the p(r) curve (Fig. 3c). The Brpt-lectin construct showed a typical pair distance distribution for an elongated protein with a globular domain at the end, with a D_max_ value of 249 Å, compared to 170 Å for the Brpt1.5 construct alone (Fig. 3c). In order to model the solution structure of the lectin-Brpt construct, 50,000 models were generated using SASSIE based on the lectin domain structure and previous structures of other Brpt1.5 constructs ^26, 35^ as described ^43^, allowing flexibility at the hinge between the lectin domain and the start of the B-repeat domains as well as flexion within the B-repeat region. After removal of models with clashes, the 12,000 remaining models were tested for consistency with the observed scattering curves using EOM (Fig. 3d-e). The best-fit models indicate that lectin domain adopts a moderately flexible position at the start of the B-repeat region (furthest from the staphylococcal cell wall), with the Y580 putative binding pocket ideally positioned to interact with a corneocyte receptor (Fig. 3f-g, Fig. S3). Interestingly, this arrangement is reminiscent of chaperone-usher pili from gramnegative bacteria, with a highly extended rod tipped by a lectin subunit for host adhesion ^53–55^.

**Fig. 3:**
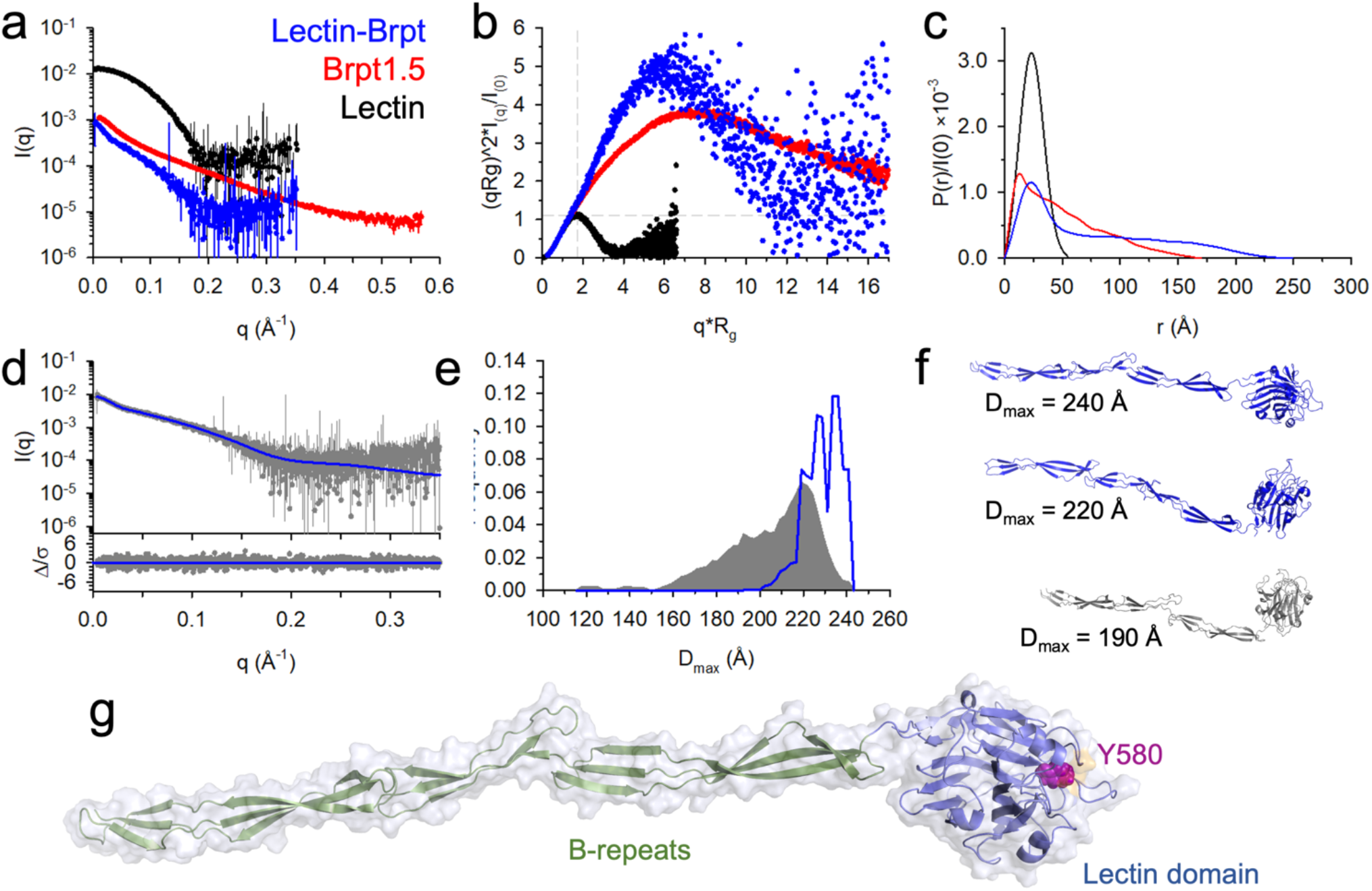
SAXS solution structural characterization of Aap-Brpt. (**a**) The SAXS scattering curves for Lectin (black), Brpt1.5 (red), and Lectin-Brpt (blue). (**b**) The dimensionless Kratky plot. Dashed lines indicate the intersection at which a Gaussian peak should be observed for a globular particle. (**c**) P(r) distributions demonstrating overall particle dimensions, where D_max_ is determined by the intersection with the x-axis. (**d**) The EOM fit (blue line) of the experimental data (gray), with the difference plot in the lower panel. (**e**) The D_max_ distribution from EOM showing the distribution of SASSIE-generated models (gray) based on flexibility of the hinge region between the lectin and B-repeat, as well as flexion along the Brpt1.5 region. The distribution of the best-fit EOM models is shown in blue. (**f**) The top two models (blue) were frequently chosen as part of the EOM ensemble, while the lower model (gray) is from the modeling pool, but not in the range of D_max_ of the ensemble. (**g**) Structure of the most frequently chosen model from the EOM ensemble, highlighting the position of the Y580 binding pocket.

However, such pili are formed by the concerted assembly of 5 or more major and minor pilin subunits and translocated by a dedicated membrane usher, whereas Aap acts as a single-molecule, multifunctional pilus-like structure.

The distal end of the lectin domain as it sits on the highly elongated B-repeat superdomain contains a series of loops; three of these are unusually long compared to other legume lectins or SraP, the closest structural homolog to Aap lectin (Fig. 1c, 1f). Two of the long loops (the 380 and 520 loops) are on the back side of the lectin domain, whereas the 580 loop originates from the Y580 binding pocket (Fig. 1c, 2a). These three loops help to form a broad platform near the distal end of the lectin domain that is likely involved in ligand recognition by Aap (Fig. 1f). To better understand the structural dynamics of these long distal loops as well as the Ca^2+^-binding and Na^+^-binding loops, we conducted multidimensional NMR studies on Aap lectin. Several key residues in and near the Y580 loop could not be directly visualized, presumably because of intermediate conformational exchange; R1, R2, and hetNOE relaxation measurements confirmed that the same regions surrounding Y580 showed a high degree of conformational exchange (Fig. 4, Fig. S4). Conformational exchange in these loops is consistent with a pocket that is able to bind to several different glycans.

**Fig. 4:**
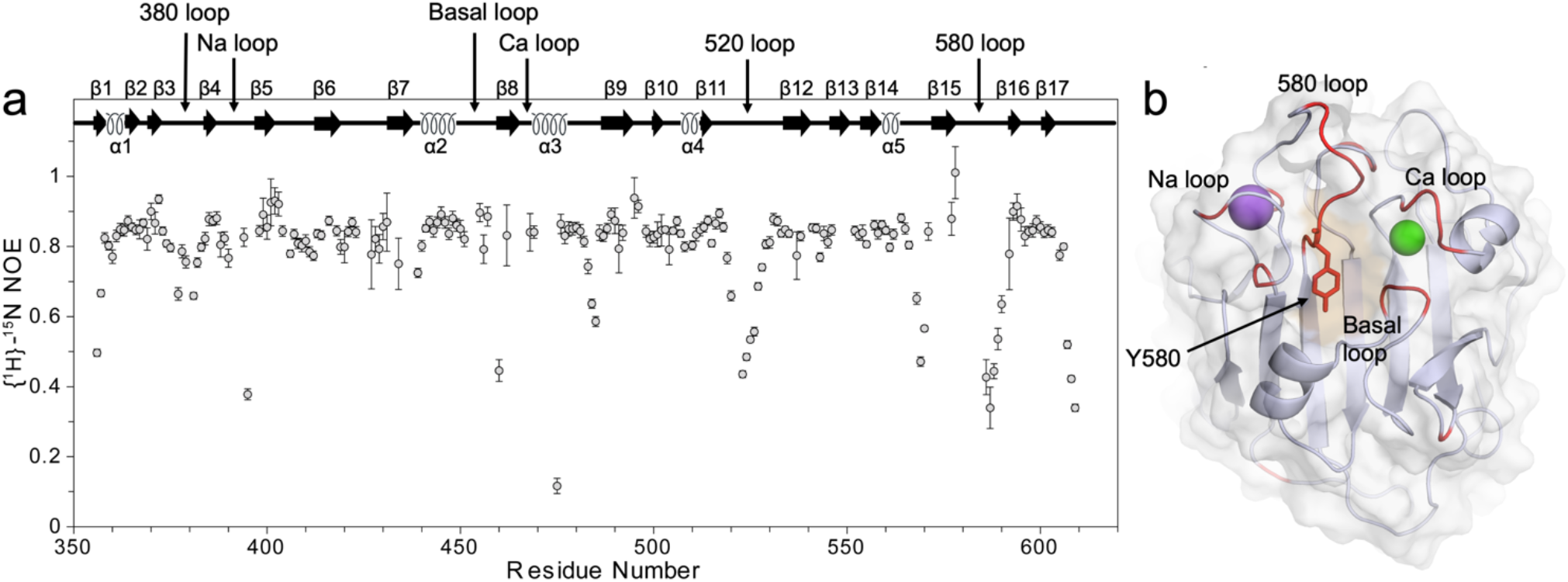
NMR reveals conformational exchange of Aap lectin loops. (**a**) {^1^H}-^15^N heteronuclear NOEs (hetNOE) of Aap-Lectin indicate the flexibility of loops surrounding the glycan binding site. (**b**) The residues that could not be assigned in the solution NMR study of Aap-lectin are highlighted in red. Residues near the Y580 pocket include Q579 through S585 in the 580 loop; N471 through D473 in the Ca loop; L452-E454 in the basal loop; and residues E391, T396, and T397 in the Na loop.

### Glycan specificity of the Aap lectin domain

In order to define the glycan specificity, Aap lectin domain was submitted for a glycan array screen (Fig. 5a). The ten highest-affinity hits are illustrated in Fig. 5b. The majority of the hits were complex biantennary or triantennary N-linked glycans containing a repeating pattern of alternating Gal-GlcNAc residues (I,e, poly-N-acetyllactosamine) at the terminal ends of the branching structures, with occasional decoration of the terminal Gal with sialic acid. Other hits included linear or branched glycans that again showed the N-acetyllactosamine repeats, occasionally with fucosylation of the GlcNAc residues. The pattern of these top hits from the glycan array are consistent with data from Roy et al. who showed that Aap lectin-mediated adhesion of *S. epidermidis* to human corneocytes was inhibited by enzymatic degradation by galactosidase, neuraminidase, and fucosidase, which cleave galactose, sialic acid, and fucose residues, respectively ^34^. To confirm these data, we conducted isothermal titration calorimetry binding experiments using N-acetyllactosamine and 3’-sialyl-N-acetyllactosamine (sialyllactosamine). ITC data showed binding of N-acetyllactosamine with ~130 μM affinity, whereas siallylactosamine bound with 10-fold higher affinity (K_D_ = 13 μM) (Fig. 5c). As described above, the putative binding site on Aap is a pocket with the aromatic ring of Y580 centered at the base of the cavity. A common recognition mechanism in lectins involves the van der Waals packing of the nonpolar face of a glycan ring against an aromatic lectin residue, accompanied by H-bonds formed between lectin residues and the axial and equatorial hydroxyl groups from the sugar ring ^56, 57^. Thus, Y580 is a likely candidate for a stacking interaction with a bound glycan. We produced a Y580A mutant of Aap lectin and determined that the mutant protein failed to bind either N-acetyllactosamine or siallylactosamine (Fig. 5d). Despite repeated attempts, neither lactosamine-based glycan could be co-crystallized with Aap due to crystal packing interactions that sterically blocked the binding site. In addition, NMR titrations consistently showed no chemical shift perturbations in the presence of 20 mM lactosamine (Fig. S5). The relatively low affinity for N-acetyllactosamine may be partly due to the flexibility in the region of the binding site detected by NMR (Fig. 4); however, the affinities observed for both glycan ligands are within the range reported for N-acetyllactosamine or siallylactosamine-containing N-glycans binding to legume lectins ^58, 59^.

**Fig. 5:**
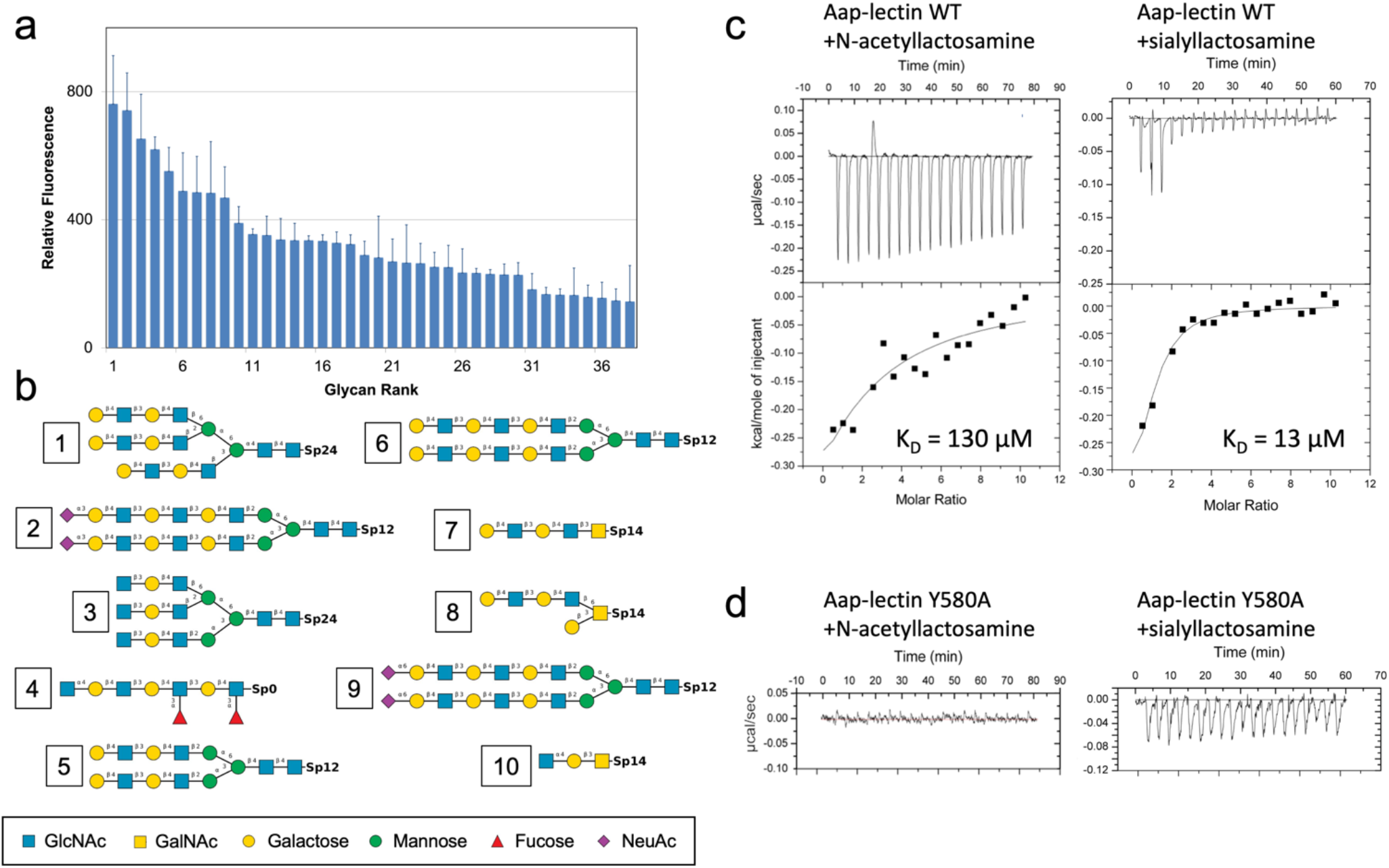
Glycan specificity of Aap lectin. (**a**) Relative fluorescence measurements for labeled Aap lectin domain binding to a glycan array. (**b**) Structures of the top 10 glycan hits from the array. Note that all top hits share a repeating Gal-GlcNAc disaccharide (N-acetyllactosamine) that can be modified by sialylation at the terminal end. (**c**) ITC data for Aap lectin binding to N-acetyllactosamine and sialyllactosamine. **(d**) ITC data for Aap-Y580A mutant binding to N-acetyllactosamine and sialyllactosamine.

### *S. epidermidis* adhesion to human corneocytes involves glycan binding via the Y580 pocket

To validate our structure-guided mutagenesis and binding data, we measured adhesion of a stably GFP-expressing *S. epidermidis* 1457 Δ*ica* strain to healthy human corneocytes sampled from healthy volunteers by tape stripping. The Δ*ica* strain of *S. epidermidis* has had the *icaADBC* operon deleted, which is responsible for the biosynthesis of the PNAG (PIA) biofilm polysaccharide; this strain was used to preclude PNAG-mediated adhesion. The *S. epidermidis* 1457 Δ*ica* strain adhered well to corneocytes in contrast to the Δ*icaΔaap* negative control (Fig. 6a). Consistent with the glycan-binding function of the lectin domain, addition of soluble N-acetyllactosamine inhibited *S. epidermidis* adhesion at all concentrations tested (ranging from 62.5–1000 μM) (Fig. 6a); likewise, soluble sialyllactosamine inhibited adhesion at ten-fold lower concentrations (12.5–100 μM; 6.25 μM showed a trend toward inhibition but did not reach significance) (Fig. 6b). These inhibition data are consistent with the ITC results showing 10-fold higher binding affinity for sialyllactosamine (Fig. 5c). *S. epidermidis* adhesion was also inhibited by the addition of soluble recombinant Aap lectin domain at 5 μM, whereas the non-binding Y580A mutant of Aap lectin completely failed to inhibit adhesion of *S. epidermidis* 1457 Δ*ica* (Fig. 6c), confirming the role of the Y580 binding pocket of Aap lectin in skin colonization by *S. epidermidis*.

**Fig. 6:**
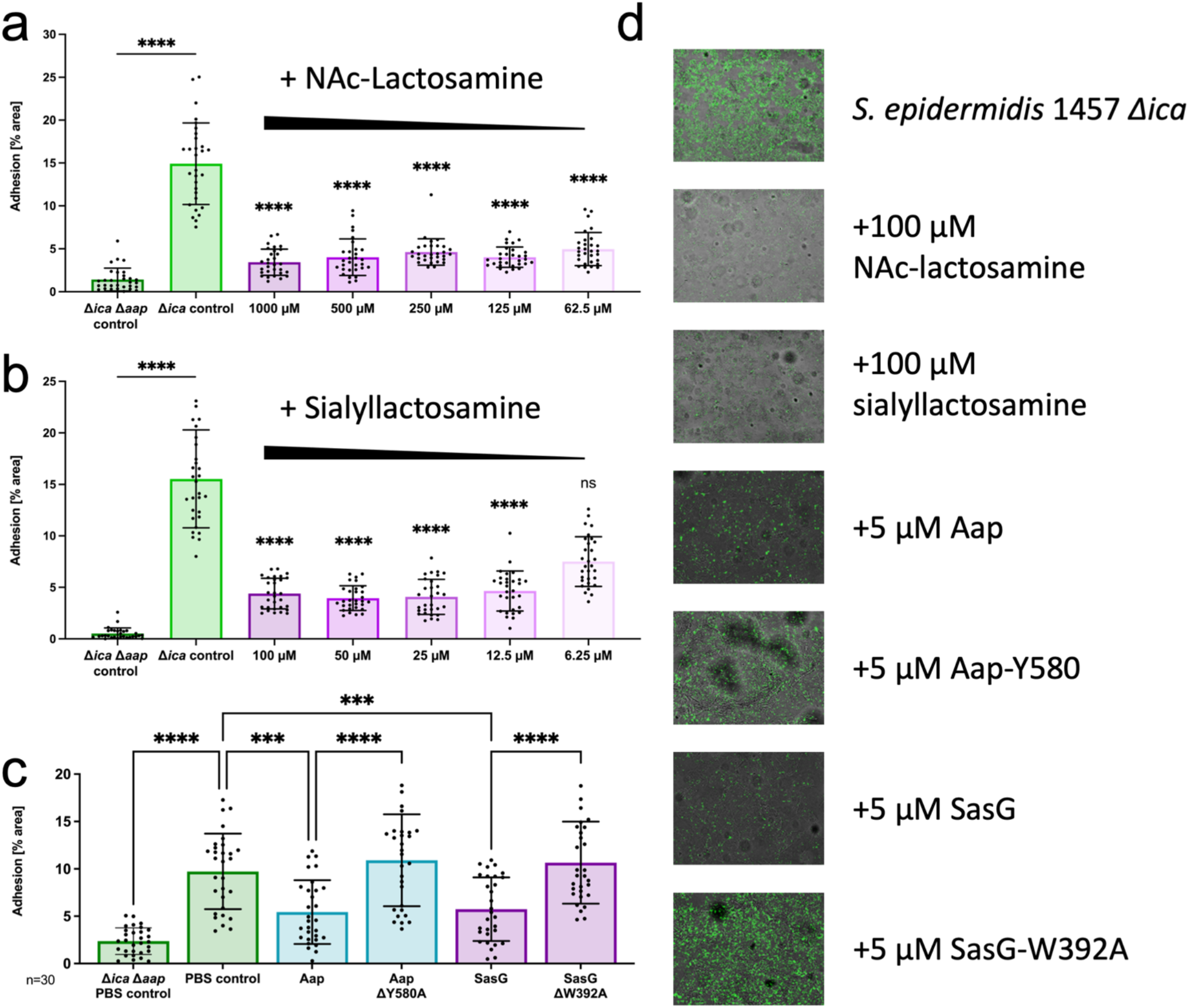
Corneocyte adhesion by *S. epidermidis* is mediated by Aap-glycan interactions. (**a**) Adhesion of *S. epidermidis* 1457 *Δica* compared to the negative control *S. epidermidis* 1457 *ΔaapΔica*, both stably expressing GFP. The *Δica* background removes production of PNAG (PIA), a polysaccharide involved in intercellular adhesion. N-acetyllactosamine inhibited *S. epidermidis* adhesion at all concentrations tested. (**b**) Sialyllactosamine inhibited adhesion of *S. epidermidis* at all concentrations except the lowest (6.25 μM). Note that sialyllactosamine was tested at 10-fold lower concentrations than N-acetyllactosamine. (**c**) *S. epidermidis* adhesion was inhibited by the addition of 5 μM soluble recombinant Aap lectin or recombinant SasG lectin, but inhibition was reduced significantly upon addition of Aap-Y580 or SasG-W392 mutant lectins. (**d**) Representative microscopy images used for the quantitative results.

### Single-cell force spectroscopy reveals tight adhesion of *S. epidermidis* mediated by Aap

We quantified the strength of the interaction between the Aap A domain and human corneocytes using AFM-based single-cell force spectroscopy ^60, 61^. Living *S. epidermidis* cells expressing variants of cellsurface Aap were immobilized on AFM cantilevers and used to image and force probe corneocytes in the quantitative imaging (QI) mode ^62^. The results (Fig. 7a) demonstrated that *S. epidermidis* 1457 *ΔsepAΔica* (expressing full-length Aap onto its surface) engages in very strong interactions with corneocytes, with adhesion forces up to 3.5 nN. Adhesive events were densely and homogeneously distributed over the corneocyte surface, with an approximate binding probability of 90% (estimated from the QI map presented in Fig. 7a). In contrast, both *S. epidermidis* 1457 *ΔaapΔica* (lacking Aap) and *S. epidermidis* 1457 *ΔsarAΔica* (lacking the A domain) showed minimal adhesion, providing direct evidence that the interaction is mediated by the Aap A domain.

**Fig. 7:**
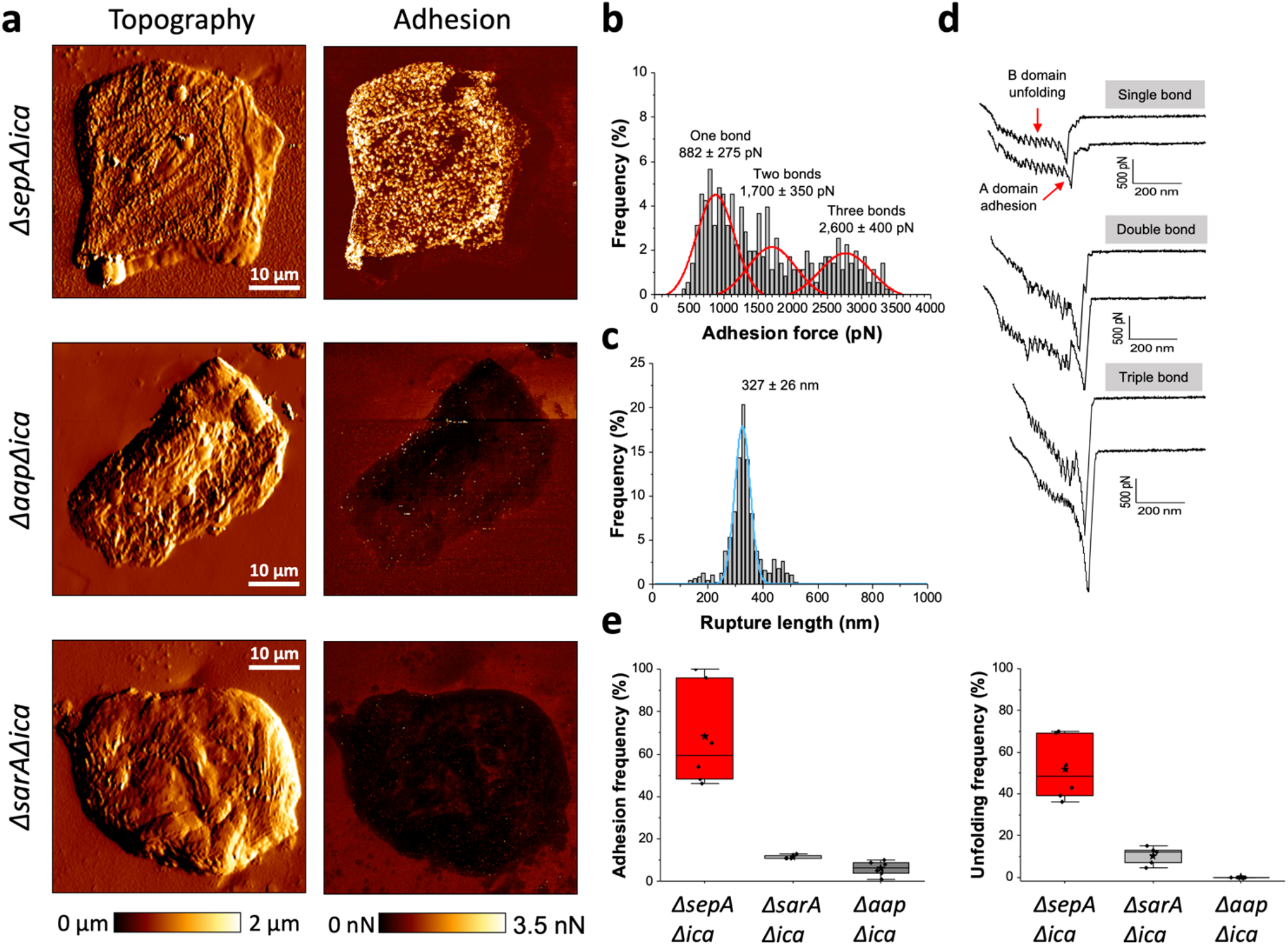
Nanoscale analysis of Aap binding to human corneocytes. **(a)** High resolution topography (left) and corresponding adhesion (right) images of human corneocytes recorded in PBS, in the quantitative imaging mode, using (from top to bottom) single *S. epidermidis ΔsepAΔica, ΔaapΔica* and *ΔsarAΔica* probes. *S. epidermidis* 1457 *ΔsepAΔica* lacks the protease SepA that cleaves the A domain from Aap, and the *Δica* background removes PNAG production; *S. epidermidis* 1457 *ΔsarAΔica* lacks the SarA global transcriptional regulator that suppresses SepA expression, resulting in SepA overexpression and thus complete proteolytic removal of the A domain of Aap. The data shown are representative from a total of 2 different cell-corneocyte pairs for each strain, including independent cultures and preparations. **(b, c)** Histograms of adhesion forces (**b**) and rupture lengths (**c**) of the *S. epidermidis* 1457 *ΔsepAΔica* - corneocyte interaction. Histograms present data merged from four different cell-pairs probed in the force volume mode. Maxima were fitted by a Gaussian model to extract mean values (± s.d.). **(d)** Representative adhesive curves documenting rupture of single, double, and triple bonds. The data in **b-d** are representative from a total of 6 different cell-corneocyte pairs, including independent cultures and preparations. **(e)** Box plots showing the binding probability (left) and unfolding frequency (right) of the *S. epidermidis* 1457 *ΔsepAΔica (n =* 6), *S. epidermidis* 1457 *ΔsarAΔica (n =* 5) and *S. epidermidis* 1457 *ΔaapΔica (n =* 5). Stars are the mean values, lines the medians, boxes the 25-75% quartiles and whiskers the s.d.

Further dissection of the adhesive interactions between *S. epidermidis* 1457 *ΔsepAΔica* and corneocytes by means of AFM force volume (FV) analysis (Fig. 7b-c and Fig. S6; total of 1000 adhesive curves from four different cell pairs) showed a large distribution of adhesion forces, ranging from 500 pN to 3,500 pN. These adhesive events were associated with a rupture length centered around 330 nm, matching the length of fully stretched adhesins, as previously demonstrated with Aap homophilic interactions involving the extension of only one Aap adhesin ^29^ (Fig. 7c). Gaussian fits of the force distributions revealed three maxima (at ~900, 1,700 and 2,600 pN) that correspond to single, double, and triple bonds (Fig. 7b). Hence, bacteria engage in multiple strong bonds with their corneocyte ligand(s), and we expect that multivalency will strengthen bacterial adhesion. Retraction force profiles (Fig. 7d) featured sawtooth patterns with multiple equally spaced peaks originating from the unfolding of individual E and G5 repeats of the B region, followed by a single strong peak resulting from the rupture of the A domain-corneocyte interaction. Conversely, the *S. epidermidis* 1457 *ΔaapΔica* and 1457 *ΔsarAΔica* strains showed only few adhesion events coupled with small forces and the curves associated presented slight to no unfolding pattern (Fig. 7e and Fig. S7a,b). These results support even more the key role of the Aap A domain in this interaction. Excitingly, the 900 pN bond strength (obtained using the *S. epidermidis* 1457 *ΔsepAΔica* strain) is an order of magnitude higher than that of single lectin-carbohydrate interactions ^63^, implying that the A domain does not simply bind through a single “classical” lectin interaction. It is also larger than the strength recently reported for single Aap-Aap bonds (~600 pN) ^29^. The G5-E repeats unfold at mechanical forces larger than the unfolding forces of β-fold multidomain proteins (<200 pN), indicating they are highly stable. Under tensile loading, the repeats will act as force buffers capable of relieving mechanical stress. Folded sacrificial domains in *S. epidermidis* adhesive polyproteins have been shown to maximize mechanical work dissipation and were suggested to serve as an adhesion strategy by bacteria ^64^. The B-repeat superdomain may play a role in strengthening of the lectin-domain mediated interaction. The unusual mechanostability of the A domain-ligand complexes and of the B repeats is of biological relevance as it allows the pathogen to tightly attach to the skin under mechanical stresses occurring in vivo (e.g., shear, epithelial turnover, cell surface contacts).

### SasG lectin adopts a similar fold and glycan specificity

Similar to *S. epidermidis, S. aureus* expresses a large CWA protein named SasG with the same overall domain architecture as Aap that shares many of the same functions (Fig. S1) ^46^. Specifically, its A domain is implicated in adhesion to host tissue such as desquamated nasal epithelial cells ^46–48^ and the B-repeat superdomain engages in homophilic (or heterophilic) assembly interactions with other SasG (or Aap) molecules in a Zn^2+^-dependent manner ^44, 49^. We solved the crystal structure of the SasG lectin to 1.93 Å resolution, revealing the same basic fold as Aap (Fig. 8a). The structural Ca^2+^ ion is bound in the corresponding location to the site in Aap, and the Ca^2+^-binding loop adopts nearly an identical configuration in the two lectins; similarly, the basal loop-helix ‘plug’ forming the bottom boundary of the binding site is conserved (Fig. 8b). However, the SasG structure does not show a bound structural Na^+^ ion, resulting in alternate conformations of the loops corresponding to the Aap Na^+^-binding loop and 580 loop (referred to as the 390 loop in SasG). In place of Y580, SasG features a different aromatic residue, W392, in approximately the same position, which could still permit stacking of the hydrophobic face of a sugar ring with the planar aromatic rings of the tryptophan side chain. As in Aap, the conserved central aspartate (D240) is in the atypical *trans* backbone configuration, and its carbonyl oxygen forms a water-mediated interaction with the Ca^2+^ ion, while its side chain forms an H-bond with the amide nitrogen of W392. Thus, even though the Na^+^ ion is not present, the structural Ca^2+^ ion and the residues in its coordination shell add constraints that stabilize the atypical configuration of the loop containing W392 (390 loop), forming the base and upper boundary of the presumed binding pocket. We confirmed that SasG lectin is able to bind N-acetyllactosamine by ITC (Fig. 8c) but binding is abrogated in the W392A mutant (Fig. 8d), confirming that this binding pocket supports binding of lactosamine, as seen for Aap. Interestingly, SasG forms a much smaller binding pocket than Aap, raising the possibility that either N-acetyllactosamine binding involves an induced fit conformational change or that Na^+^ binding, not observed in the SasG structure, may be needed to provide additional constraints that hold the binding pocket in the open state.

**Fig. 8:**
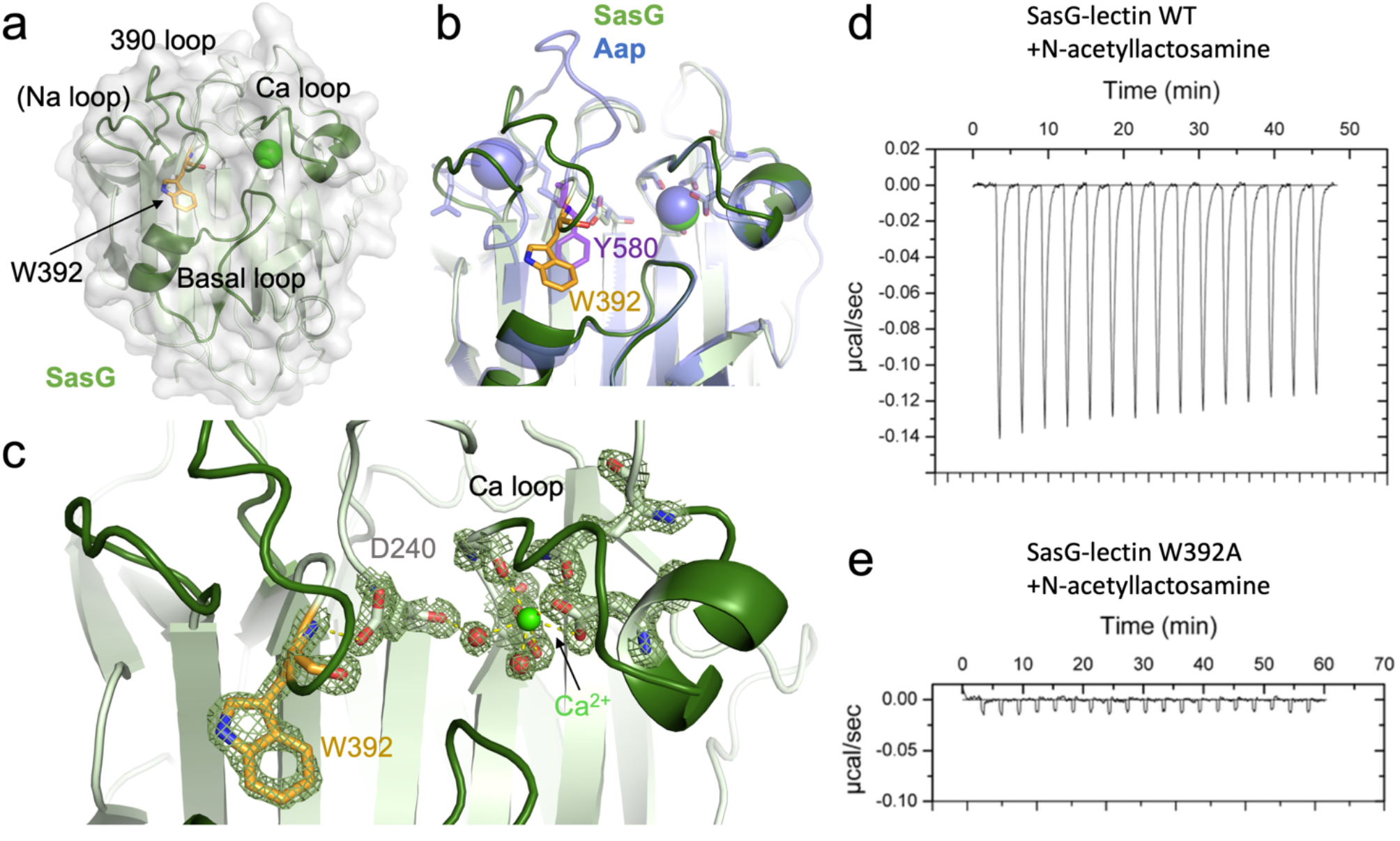
SasG lectin adopts a similar structure and binding specificity to Aap. (**a**) Crystal structure of the *S. aureus* SasG lectin domain illustrated with a transparent surface, showing the loops that converge on the front face of the lectin to form the binding pocket. (**b**) Superposition of SasG and Aap lectin domains. Residues W392 (SasG) and Y580 (Aap) that form the floor of each binding pocket are shown as sticks. Note that SasG features the structural Ca^2+^ ion but does not contain the structural Na^+^ ion observed in Aap. (**c**) Details of the interaction network involving Ca^2+^, conserved aspartate D240 (which forms a *trans* peptide bond, similar to D431 in Aap), and W392, all of which stabilize the sharp bend in the main chain near W392 that creates the floor and upper wall of the binding pocket. (**d**) ITC data for SasG binding to N-acetyllactosamine; the binding was weak and could not be satisfactorily fitted to a binding model. (**e**) ITC data for SasG-W392A mutant binding to N-acetyllactosamine.

### SasG mediates staphylococcal adhesion to human corneocytes via glycan binding to the W392 pocket

To test the role of SasG in staphylococcal adhesion to human corneocytes, we conducted corneocyte adhesion assays using the non-adherent coagulase-negative species *S. carnosus* engineered to express SasG on its surface, *S. carnosus*/pALC2073-*sasG*_COL_ (*S. carnosus-sasG*), versus an empty-vector negative control. This approach avoided complications of wild-type *S. aureus* gene regulation and other potentially competing surface adhesins. The *S. carnosus-sasG* strain adhered well to healthy human corneocytes whereas the empty-vector control failed to adhere (Fig. 9a). As observed for *S. epidermidis*, N-acetyllactosamine at concentrations ranging from 62.5 to 1000 μM inhibited adhesion of *S. carnosus-sasG*, as did sialyllactosamine at ten-fold lower concentrations (12.5–100 μM) (Fig. 9a-b). Furthermore, soluble SasG lectin at 5 μM inhibited adhesion of *S. carnosus-sasG*, but the W392A mutant of SasG lectin inhibited adhesion to a significantly lesser degree than wild-type lectin (Fig. 9c). These data confirm the role of the SasG lectin domain in skin colonization and the specific role of the W392 glycan binding site.

**Fig. 9:**
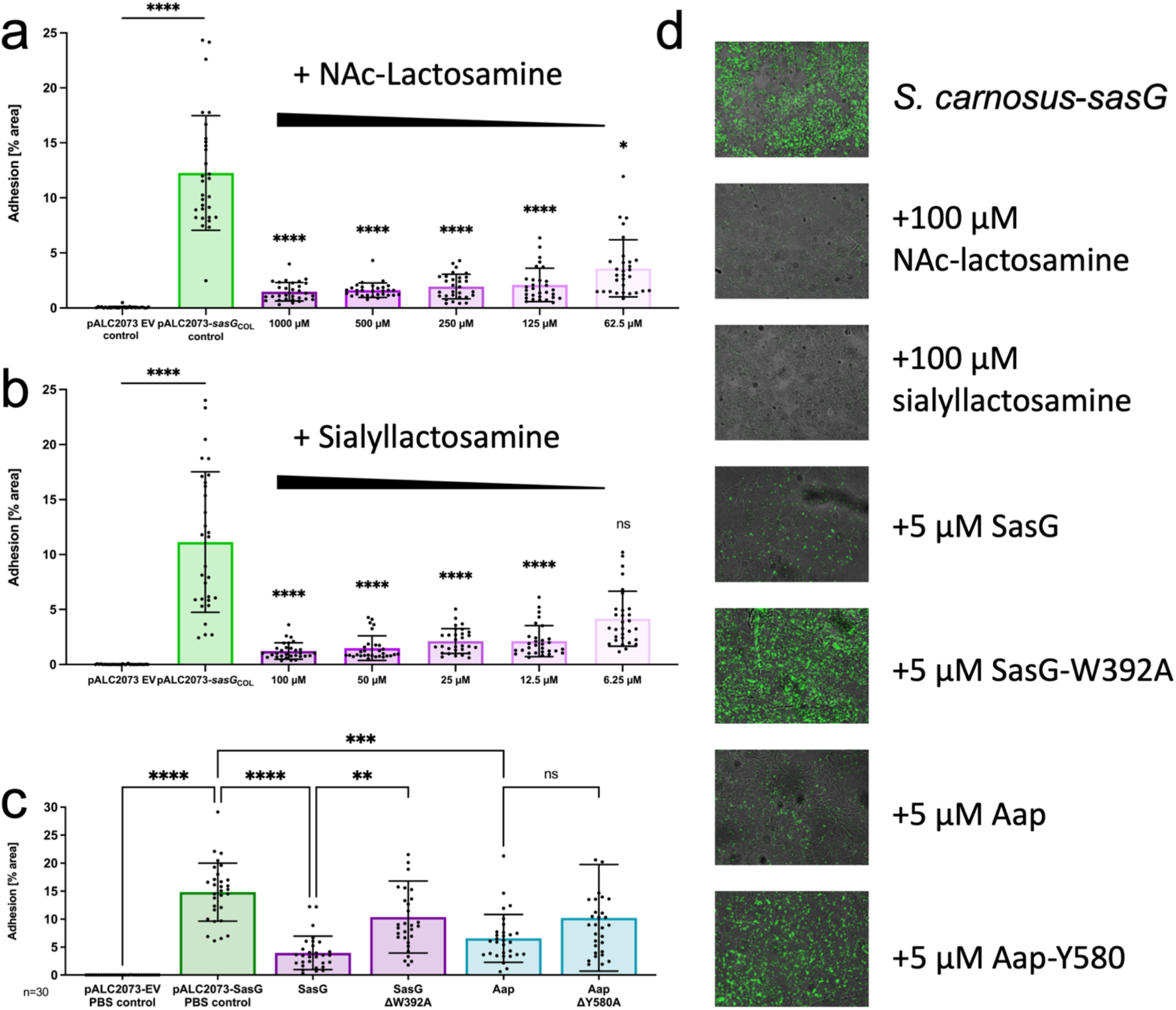
Staphylococcal adhesion to corneocytes mediated by SasG. (a) Adhesion of *S. carnosus* engineered to express SasG, *S. carnosus/pALC2073-sasG*_COL_ (*S. carnosus-sasG*), compared to the empty-vector negative control. N-acetyllactosamine inhibited *S. carnosus-sasG* adhesion at all concentrations tested. (b) Sialyllactosamine inhibited adhesion of *S. carnosus-sasG* at all concentrations except the lowest (6.25 μM). Note that sialyllactosamine was tested at 10-fold lower concentrations than N-acetyllactosamine. (c) *S. carnosus-sasG* adhesion was inhibited by the addition of 5 μM soluble recombinant SasG lectin but inhibition was reduced significantly upon addition of SasG-W392 mutant lectin. Soluble Aap lectin also inhibited adhesion; addition of Aap-Y580 mutant showed a trend toward reduced inhibition that did not reach significance. (**d**) Representative microscopy images used for the quantitative results.

### Lectin-mediated cross-inhibition of *S. aureus* and *S. epidermidis* adhesion to human corneocytes

Given the similarities between Aap and SasG lectins in their overall structure, glycan specificity, and binding site architecture containing a prominent aromatic residue, the question arises as to whether *S. aureus* and *S. epidermidis* can compete with one another for adhesion to human corneocytes. We tested each soluble lectin for cross-inhibition of corneocyte adhesion (i.e., soluble SasG inhibiting *S. epidermidis* or soluble Aap inhibiting *S. carnosus-sasG*). We observed that 5 μM soluble SasG lectin inhibits adhesion of *S. epidermidis* to a similar degree as soluble Aap, but that the W392A mutant failed to inhibit adhesion (Fig. 6c). Likewise, adhesion of *S. carnosus-sasG* was inhibited by the addition of 5 μM soluble Aap lectin, whereas the Y580 mutant showed a clear trend toward decreased inhibition that did not reach significance; this loss of inhibitory effect was comparable to that seen with the W392A mutant of SasG lectin (Fig. 9c).

## Discussion

The data presented here demonstrate that Aap lectin domain adopts a fold generally similar to legume lectins as well as the staphylococcal CWA protein SraP ^50^, but with an atypical binding site architecture due to the central *trans* aspartate residue (D431) that interacts with the amide nitrogen of Y580, forming the floor of the glycan binding pocket. Furthermore, Aap lectin features a structural Na^+^ ion that provides additional constraints on the configuration of the glycan binding site. NMR relaxation data revealed a surprising degree of conformational flexibility in regions near the Aap binding site, particularly in the 580 loop that forms part of the glycan binding site (i.e., residue Y580) and extends from Y580 upward into one of three unusually long loops in Aap that diverge from other family members, including SraP. In addition to the long 580 loop, two other long loops extend away from the body of the lectin domain on the upper rear surface of the domain, forming a platform with significant surface area that is unique to Aap. We propose that this surface is likely to represent a binding site for the protein moiety of a corneocyte glycoprotein receptor (Fig. 10a). High-resolution AFM-based imaging showed that the corneocyte receptor recognized by Aap is expressed at high density and is distributed homogeneously across the corneocyte surface (Fig 8a). Single-cell force spectroscopy data revealed surprisingly strong adhesion forces (900 pN) for individual interactions between Aap and the corneocyte receptor; such an adhesion force is an order of magnitude stronger than that expected for typical glycan-lectin interactions. These data are consistent with a bipartite interaction between Aap lectin domain and both protein and N-glycan moieties of a corneocyte receptor rather than a simple lectin-glycan interaction alone. Previous force spectroscopy studies have revealed the remarkably high mechanostability of the B-repeat regions of both Aap and SasG ^29, 44^. Given the direct connection between the lectin domain and the start of the B-repeat superdomain (Fig. 3), we speculate that the unique mechanical properties of the unusual B-repeat protein fold ^26, 36^ may also play a role in strengthening the Aap-corneocyte interactions.

**Fig. 10:**
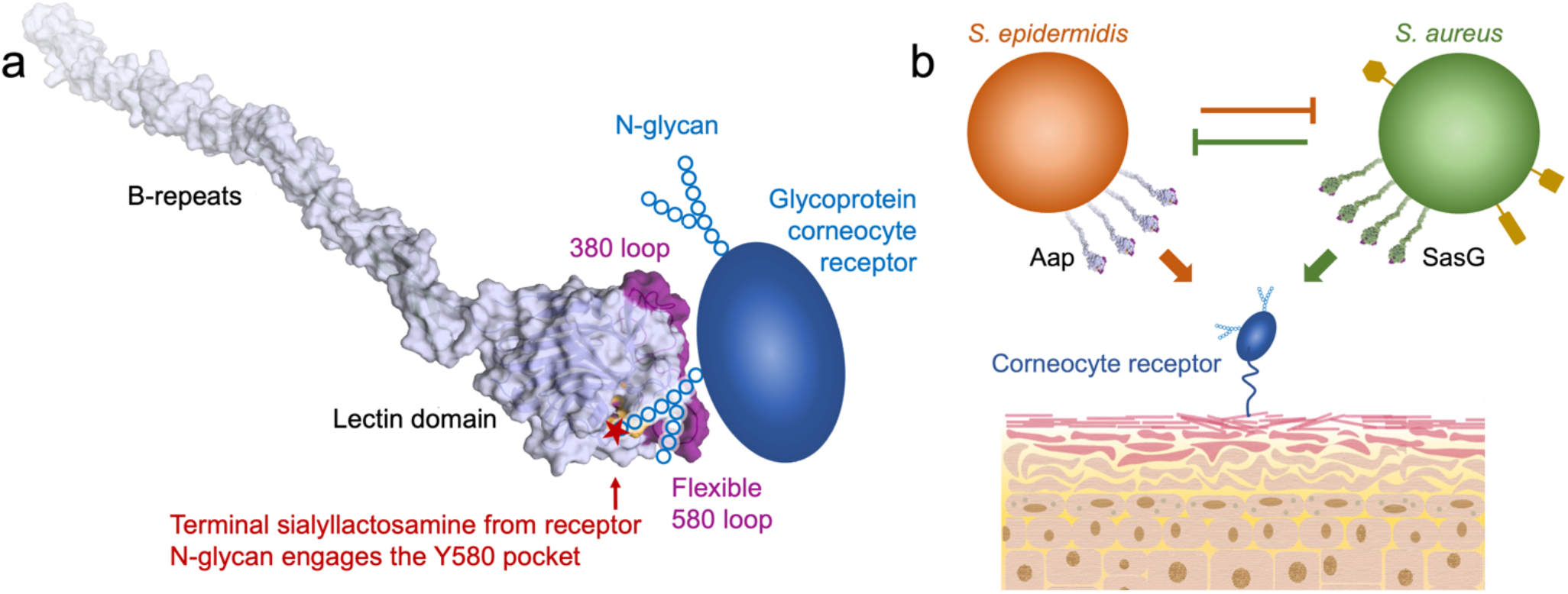
Model for corneocyte adhesion mediated by Aap and SasG. (**a**) Model for the interaction of Aap with a corneocyte glycoprotein receptor. The best-fit structural model of Aap lectin-Brpt from the SAXS EOM analysis is illustrated with a semi-transparent surface. The Y580 binding pocket is colored orange, and the divergent long loops at the top of the lectin domain are colored purple. The cartoon of the glycoprotein receptor illustrates the proposed bipartite binding mechanism in which the divergent long loops (purple) of Aap recognize the protein moiety of the receptor, and a receptor N-glycan containing terminal sialyllactosamine interacts with the Y580 binding pocket. (**b**) Schematic illustrating competition between *S. epidermidis* and *S. aureus* for corneocyte adhesion and skin colonization. Both Aap and SasG recognize the same (or highly similar) corneocyte receptor(s). Structures illustrated in yellow on *S. aureus* represent additional surface adhesins (e.g., SdrD, FnBP-B, or ClfB) with specificity for corneocyte or keratinocyte receptors that could confer an advantage to *S. aureus* under appropriate conditions.

The structure of SasG lectin is highly similar to Aap lectin, with the same atypical architecture resulting from the *trans* aspartate residue (D240) that interacts with the amide nitrogen of W392, placing an aromatic residue similar to Y580 at the base of the binding pocket. The binding pocket is notably smaller in the SasG structure compared to Aap, but this may be due to the lack of a bound structural Na^+^ ion that was observed to aid in forming the binding pocket architecture in Aap. Furthermore, the three long loops observed in Aap that diverge from SraP are also present in SasG, suggestive of similar bipartite recognition of a glycoprotein receptor and consistent with the overlapping specificity for corneocyte adhesion between SasG and Aap.

Taken together, our data strongly suggest that Aap and SasG lectin domains share specificity for a corneocyte receptor, presumably a glycoprotein, that interacts with a large surface patch on the top of the lectin domain formed primarily by the three long divergent loops (Fig. 1c, 1f, 10a). Interestingly, one of these long loops (the 580 loop in Aap), which originates in the Y580 glycan-binding pocket, shows a high degree of conformational exchange by NMR relaxation measurements (Fig. 4); the other two loops (380 loop and 520 loop) on the back side of the lectin also show increased dynamics, albeit to a lesser degree than the 580 loop. If, as expected, the protein moiety of the corneocyte receptor interacts with the distal (top) surface of the lectin, that interface on Aap would likely involve the 380, 520, and/or 580 loops (Fig. 10a). Our glycan array data suggest that the corneocyte receptor features an N-glycan decorated with 3’ sialyllactosamine at the terminal end that binds in the Y580 binding pocket. Conformational exchange in the 580 loop may be necessary to facilitate binding of terminal sialyllactosamine groups (Fig. 10a), given the variability observed for N-glycoforms on individual glycoproteins ^65–68^. The proposed contribution of both protein and glycan components of the corneocyte receptor to the binding free energy is consistent with the unusually strong adhesion force measured by single-cell force spectroscopy (Fig. 7).

The corneocyte adhesion assays (Figs 6 and 9) provided valuable insights into inter-species competition for colonizing the skin niche. Soluble Aap lectin domain inhibited corneocyte adhesion not only by *S. epidermidis*, but also by *S. carnosus-sasG;* likewise, soluble SasG lectin domain inhibited *S. epidermidis* adhesion. These data indicate that both *S. epidermidis* and *S. aureus* directly compete for the same corneocyte receptor. Both lectins were inhibited by N-acetyllactosamine and sialyllactosamine with similar dose responses. Given the near-ubiquitous colonization of skin by *S epidermidis, S. aureus* would need to out-compete *S. epidermidis* for interaction with the same (or a highly similar) corneocyte receptor (Fig. 10b). However, *S. aureus* is known to express other CWA adhesins with specificity for corneocyte or keratinocyte receptors; co-expression of SasG with one or more of these additional adhesins may provide a critical advantage when competing with established *S. epidermidis* for skin colonization (Fig. 10b). One such *S. aureus* adhesin is the CWA protein SdrD, a member of the serine/aspartate-rich family of proteins. SdrD specifically interacts with desmoglein-1 ^69^, a corneodesmosomal cadherin that is expressed at high levels in outer epidermal layers. Furthermore, fibronectin-binding protein B (FnBP-B) and ClfB have been shown to interact with loricrin, another characteristic skin-resident protein ^70, 71^, and both FnBP-B and clumping factor B (ClfB) bind to corneodesmosin ^72^. Context-dependent expression of various adhesins with specificity for skin receptors is likely to play an important role in colonization by *S. aureus* under specific conditions, such as the overgrowth of *S. aureus* during AD flares. Upregulation of particular staphylococcal adhesins or combinations thereof could recognize corneocyte receptors that are differentially expressed (or exposed) under disease conditions. For example, AD skin is characterized by decreased expression levels of the critical structural protein filaggrin, resulting in changes to the architecture of the skin layers and potential exposure of keratinocyte receptors that would be inaccessible in healthy skin. Indeed, FnBP-B and ClfB have been shown to interact with corneodesmosin that is exposed in AD skin as a result of a lack of natural moisturizing factor ^72, 73^.

The corneocyte adhesion assays also validated the critical role of the Y580 or W392 binding pockets in adhesion via Aap or SasG, demonstrating that small soluble compounds (e.g., N-acetyllactosamine or sialyllactosamine) are efficacious at inhibiting staphylococcal colonization. This opens the door to the development of small-molecule therapeutic agents that target these binding pockets on Aap or particularly on SasG and specifically inhibit adhesion to skin. *S. aureus* produces a wide range of virulence factors including proteases that further damage the skin of AD patients, decrease its barrier function, and increase inflammation, likely predisposing AD patients to develop further atopic conditions ^13, 14, 18^. Compounds targeting SasG-mediated adhesion, whether alone or in combination with other inhibitors, could find application in the case of AD or other skin conditions linked to *S. aureus*, including impetigo, cellulitis, or staphylococcal scalded skin syndrome.

## Methods

### Protein Purification

pDest-HisMBP and pDest17-His plasmids containing the Aap lectin domain (residues 351-605) from *S. epidermidis* RP62A were transformed into BLR (DE3) cells (Novagen). Single colony isolates were transferred to 100 mL of liquid media (LB) supplemented with 50 μg/ml ampicillin sodium salt. Cultures were allowed to grow for 16 hours at 37° C. The overnight (O/N) culture was diluted into 1 L flasks of LB containing 50 μg/mL ampicillin sodium salt to an initial OD600 of 0.03. Cells grew for 3-5 hours, at 37° C, until an OD600 of 0.8-1.0 was reached. Flasks were transferred to an ice bath until the culture attained a temperature of 10° C (to induce a cold-shock response), at which point 22 mLs of ethanol (to maintain cold chaperonin expression) and 300 μM Isopropyl β-D-1-thiogalactopyranoside (IPTG) were added to each flask. Cells were then allowed to grow at 18° C for an additional 16 hours. Cells were harvested via centrifugation at 5,000 RPM for 1 hr. Supernatant was discarded and the cells were resuspended in 20 mM Tris pH 7.5, 150 mM NaCl. The suspension was then frozen overnight. Cells were thawed until a homogeneous suspension was established. The cells were then sonicated in 30 mL aliquots and centrifuged at 14,000 RPM for 1 hr. Insoluble pellet fractions were discarded and the supernatant was separated from the sample. The supernatant was applied to a gravity-flow column containing NiNTA (ThermoFisher) and eluted from the resin with a gradient of 30-300 mM imidazole. Isolated Aap lectin was then dialyzed and incubated with TEV protease to remove His tag or His-tagged fusion protein (MBP). The solution was dialyzed and applied to the NiNTA to remove TEV protease and His-tagged cleaved material. Flow through was then applied to a Superdex 75 column and purified lectin fractions were pooled. Likewise, SasG lectin (residues 163-419 from *S. aureus* 502a) was expressed using the pDest-HisMBP plasmid and purified as described for Aap lectin.

### Protein Crystallization and Data Collection

All crystals were grown via the hanging drop vapor diffusion method. Protein stocks at a concentration of 20-25 mg/ml in 20 mM BisTris (pH 6.5) and 150 mM NaCl were combined with equal parts of mother liquor (1 μl +1 μL or 2μl +2 μL). Aap lectin crystals formed in various conditions: 1) 20-28% BCS PEG SMEAR Medium (Molecular Dimensions, CalibreScientific); 2) 100 mM Na Acetate (pH 4.5), 22.5% BCS PEG SMEAR High (Molecular Dimensions, CalibreScientific); and 3) 200 mM KCl, 200 mM CaCl_2_, 45% v/v 2-methyl 2-4 pentanediol and 100 mM BisTris (pH 6.5). SasG lectin crystals were grown in 100 mM Tris (pH 7.9), 40 mM Na Formate, 40 mM CaCl_2_, and 14-20% BCS PEG SMEAR Low (Molecular Dimensions, CalibreScientific). Crystals were added to a cryoprotectant consisting of 80% mother liquor and 20% MPD prior to being flash frozen in liquid nitrogen. Data collection occurred at the Advanced Photon Source at Argonne National Lab through the Northeastern Collaborative Access Team (NE-CAT), utilizing both 24-ID-C and 24-ID-E beamlines. Diffraction data were collected on Aap lectin crystals for S-SAD using an oscillation range of 0.5°, collecting 1440 frames over a resolution range of 200-1.65 Å. Native data sets used an oscillation range of 0.2°, collecting 1800 frames, over a resolution of 200-1.03 Å. SasG data sets used an oscillation range of 0.5°, collecting 360 frames, at a resolution of 200-1.93 Å.

### Phasing, Refinement, and Validation

Structural determination for Aap lectin was carried out by S-SAD phasing through the detection of three anomalous scattering atoms: sulfur atoms from two methionine residues and a bound calcium. Data was processed by the RAPD server available through NE-CAT (https://rapd.nec.aps.anl.gov/rapd); data indexing, space group assignment, scaling, and integration were carried out in RAPD via XDS ^74^. Shelx ^75^ is utilized in the RAPD suite followed by PHENIX AutoSol (HySS, PHASER, RESOLVE, Xtriage, PHENIX-Refinement) ^76^ to determine anomalous scatterers, scatterer positions, calculation of experimental phases, density modifications, model-building, and refinement. Further refinement was carried out in PHENIX, and model building was done using Coot ^77^, on local computers. The 1.06 Å native Aap lectin structure was solved by molecular replacement (MR) using Phaser-MR with the S-SAD-phased Aap as a search model. The SasG structure was solved by MR using Phaser-MR with native Aap lectin as a search model. Anisotropic refinement was carried out using PHENIX Refine. The final refined structures were validated using MolProbity ^78^. CASTp (Computed Atlas of Surface Topography of proteins) was used to calculate the volume of surface pockets, including the putative glycan binding pocket ^79^.

### Glycan array screening

Aap lectin was dialyzed into PBS pH 8.0, concentrated to 2 mg/ml, and labeled with AlexaFluor 488 (10 μl of 10 mg/ml stock solution). The reaction was allowed to incubate at 4 °C for 48 h. Labeled Aap lectin was dialyzed into BisTris pH 6.5, 150 mM NaCl to quench the labeling reaction and remove free unbound AlexaFluor. Labeled Aap at 0.5 mg/ml was sent for glycan array screening by the Center for Functional Glycomics, using array version 5.4 consisting of 585 distinct glycans. The binding buffer contained 20 mM Tris-HCl, pH 7.4, 150 mM NaCl, 2 mM CaCl_2_, 2 mM MgCl_2_, 0.05% Tween 20, and 1% BSA. Data were reported as average RFU from replicate measurements (4 values averaged from 6 original replicates, after discarding the highest and lowest values per condition) along with standard deviation and %CV (%CV = 100x standard deviation / Mean).

### Small-angle X-ray scattering

Lectin and Lectin-Brpt1.5 SEC-MALS-SAXS experiments were performed at the Advanced Proton Source (APS), Argonne, Illinois, on beam line 18ID, operated by BioCAT. Lectin samples used for SAXS were initially purified as described above. The lectin protein was dialyzed into a 50 mM MOPS (pH 7.2) and 50 mM NaCl buffer and was concentrated to 1 mg/mL prior to being transported to the BioCAT facilities. It was then concentrated on-site to 15-30 mg/mL. Samples were centrifuged at 12,000 rpm for five minutes prior to injection onto a Superose 200. Prefiltered sample aliquots of 237 μl were applied to a Superose 6 (10/300 GL) column (Cytiva) on an Agilent 1260 Infinity II HPLC at a flow rate of 0.6 ml/min. The samples then passed through the column and were measured using, in order, an Agilent UV detector, a MALLS detector, a Wyatt Technologies Dawn Helios II, and an Optilab RI detector. Intensity measurements of scattering were taken using a Dectris Pilatus3 X 1M detector. Over a time course of 1 second, samples were exposed for 0.5 seconds. Brpt1.5 SAXS data were collected at The Advanced Light Source (ALS) at Lawrence Berkeley National Laboratory using the mail-in HT-SAXS service on SIBYLS beamline 12.3.1 ^80–83^. Brpt1.5 was dialyzed into 50 mM MOPS pH 7.2, 50 mM NaCl, 2% glycerol and data collected on samples at 5 mg/mL, 6.7 mg/mL, 10 mg/mL, and 15 mg/mL.

SAXS analyses were performed in BioXTAS RAW (version 2.1.0) ^84^. The concentration series collected for Brpt1.5 was extrapolated to zero concentration in Primus from ATSAS (version 3.0.3) ^85^ and then analyzed in BioXTAS RAW alongside the Lectin and Lectin-Brpt1.5 datasets. To model flexibility of Lectin-Brpt1.5, SASSIE’s monomer Monte Carlo module ^86^ was used to generate 50,000 models. After removing clashing models, approximately 12,000 models remained. Flexible regions were allowed to sample up to 30° during each move. Flexible regions included the Lectin-Brpt1.5 hinge region (604-610) and the interfaces between each G5 and spacer domain of Brpt1.5 (686-689 and 735-738). CRYSOL ^87^ in non-fitting mode was used to generate SAXS scattering curves for each model, while also generating input files for the Ensemble Optimization Method (EOM) ^88, 89^ using a custom python script ^43, 90^. CRYSOL settings were: max order of harmonics = 99; order of Fibonacci grid = 18; number of points = 101. CRYSOL was also run in fitting mode with the same settings to obtain single-model fits of the experimental dataset.

### Nuclear magnetic resonance spectroscopy

Uniformly ^2^H,^12^C,^15^N–labeled Aap lectin was expressed and purified as described above. All NMR relaxation experiments (Heteronuclear ^91–15^N NOE, T_1_, and T_1ρ_ experiments) were collected at 298K on a Bruker AVANCE III 600 MHz spectrometer with 0.5 mM lectin in NMR buffer (20 mM HEPES, pH 6.8, 50 mM NaCl) containing 10% D2O. The NMR data was referenced for ^1^H chemical shifts by using 2, 2-dimethyl-2-silapentane-5-sulphonic acid (DSS) at 298 K as a standard. The ^13^C and ^15^N chemical shifts were referenced indirectly. The steady-state heteronuclear {^1^H}–^15^N NOE was measured as the ratios of the peak intensities with and without proton presaturation period (4 s) applied before the start of the ^1^H–^15^N TROSY HSQC experiment. The T_1_ and T_1ρ_ experiments were recorded using a pulse program described by Lakomek *et al*. ^92^. The spin-lattice T_1_ experiments were collected as pseudo-threedimensional (3D) TROSY-HSQC relaxation experiments with eight relaxation delays (0, 80, 160, 240, 320, 400, 640, and 900 ms). The spin-spin relaxation rate, *R*_2_, was determined from spin-lattice in rotating frame T_1ρ_ ^93^ collected similarly as T_1_ with eight relaxation delays (2, 12, 24, 48, 72, 96, 120, and 144 ms). The T_1ρ_ spin lock power was 2 kHz, and temperature compensation was used for both the T_1_ and T_1ρ_ experiments. Relaxation experiments were acquired in an interleaved manner with 512 × 1344 (t1 × t2) with a spectral width of 2128.62 and 9615.38 Hz. Spectra were processed using NMRPipe ^94^ program, and the relaxation rates, *R*_1_ and *R*_2_ were determined by fitting peak intensities as a function of relaxation decay time using NMRFAM-SPARKY ^95^. Titrations in the presence of 20 mM lactosamine were collected at 298 K at 600 MHz under similar conditions as described above, using 0.5 mM ^15^N labeled lectin in NMR buffer.

### Isothermal titration calorimetry

Isothermal titration calorimetry (ITC) experiments were carried out using a MicroCal VP-ITC microcalorimeter. Standard experiments were performed at 25 °C in buffer containing 50 mM phosphate (pH 7.2) and 150 mM NaCl. Briefly, the sample cell contained 20 μM of either Aap-lectin or SasG-lectin domains (1.5 mL) while the syringe had 1 mM of glycan (450 μL). Twenty injections of glycan were delivered, the first of which was 2 μl followed by nineteen injections of 14 μl each. Controls experiments were performed with 1 mM glycan in the syringe and used to correct for heat of dilution. All lectin protein concentrations were calculated using either a ThermoScientific NanoDrop or a ThermoScientific BioMate 3S UV Spectrophotometer, where the absorbance at (A280) at 280 nm was measured. Glycan concentrations were calculated based on resuspension of purchased lyophilized glycans. Fitting of all ITC data to binding models was carried out using ORIGIN software.

### Corneocyte collection

Corneocytes were collected as described in Mills et al. 2022 ^96^. Corneocytes were collected from either the lower or upper arm near the elbow from human volunteers with healthy skin. The selected skin area was cleaned with an alcohol wipe and allowed to air dry. Corneocytes were removed using clear, adhesive tape stripping discs (d-Squame D100; Clinical & Derm).

### Preparation of bacterial strains

Bacterial strains used in this study are described in Table S2. Superfolder green fluorescent protein (sGFP)-expressing bacterial cultures for corneocyte adhesion assays were prepared as described in Mills et al. 2022 ^96^. 5 mL of *S. epidermidis* and *S. carnosus* strains were grown overnight in tryptic soy broth (TSB) at a 1:5 medium-to-culture tube ratio at 37°C with shaking aeration at 220 rpm. Cultures were grown with 10 μg/mL chloramphenicol for plasmid maintenance. Cultures were then diluted 1:50 in TSB with chloramphenicol and grown to an OD600 of ~0.75. Cultures were washed once in phosphate-buffered saline (PBS) at a 1:1 ratio and diluted in PBS to a final OD600 of 0.15 (~10^7^ CFU/mL). An OD600 of 0.15 allows for sufficient cell enumeration without bacterial clumping ^34, 96^.

### Corneocyte adhesion assay with purified glycans and lectins

Purified N-Acetyl-D-lactosamine (Sigma-Aldrich) was diluted in PBS to 1000 μM. Serial dilutions were prepared at a 1:2 ratio in 300 μL PBS from 1000 μM to 62.5 μM. Purified 3’-Sialyl-N-acetyllactosamine (Sigma-Aldrich) was diluted in PBS to 100 μM. Serial dilutions were prepared at a 1:2 ratio in 300 μL PBS from 100 μM to 6.25 μM. The corneocyte adhesion assays were adapted from Mills et al. ^96^. Adhesion of *S. epidermidis Δica* (which acts as a WT control due to this locus not contributing to intercellular adhesion ^34^) and *S. carnosus-SasG*_COL_ were tested for adhesion to healthy human corneocytes after co-incubation with purified glycans. 300 μL of prepared bacterial cultures and 300 μL of individual glycan serial dilutions as described above were mixed and co-incubated at room temperature for 20 minutes. The entire 600 μL mixture was then incubated on corneocytes for 45 minutes at 37 °C. Corneocytes were incubated with 300 μL of either *S. epidermidis Δica Δaap* or *S. carnosus*-pALC2073 (EV) as negative controls.

Adhesion of *S. epidermidis Δica* (and *S. carnosus*-SasG_COL_) were also tested for adhesion to healthy human corneocytes after pre-incubation/blocking with purified lectins. Purified recombinant Aap, Aap ΔY580A, SasG, and SasG ΔW392A lectin domains were diluted in 300 μL PBS to a concentration of 5 μM. 300 μL of purified lectins were incubated on corneocytes for 20 minutes at room temperature followed by incubation with 300 μL of prepared bacteria (see above). Corneocytes were incubated with 300 μL of either *S. epidermidis Δica Δaap* or *S. carnosus*-pALC2073 (EV) as negative controls.

### Imaging and statistical analysis

Corneocytes were imaged using fluorescence microscopy at X20 magnification using the bright-field and green channels. Ten images of each disc with corneocytes from three independent experiments (n = 30) were measured for adhesion percent area with Fiji ImageJ and analyzed in GraphPad Prism as described ^96^.

### Atomic force microscopy

Stationary phase cultures of *S. epidermidis* 1457 *ΔsepAΔica*, *ΔaapΔica*, and *ΔsarAΔica* bacteria were grown overnight in TSB (+ 10 μg/ml trimethoprim for *ΔaapΔica* only) while shaking (180 rpm) at 37 °C. They were then collected by centrifugation (5’ at 2000xg), washed two times with PBS and resuspended in PBS, with a 50-fold dilution. A small amount (50 μL) of the suspension was then deposited on the left side of a polystyrene Petri dish and the cells were allowed to adhere for 10’ then washed with PBS. On the right side of the dish, a piece of corneocytes sample (5 mm x 5 mm) was immobilized using double-sided tape. The Petri dish was filled with 3 mL PBS. For preparing single-cell probes, tipless NPO-10 cantilevers (Bruker) were functionalized on their end with single glued silica microspheres (6.1 μm diameter) and incubated for 1h in a polydopamine solution. The nominal spring constant of the colloidal probe cantilever (typically ~0.06 N/m) was determined by the thermal noise method. It was then brought into contact with single *S. epidermidis* bacteria weakly adhered to the surface of the Petri dish and retracted to attach the cell. Next the probe was transferred to the corneocytes area to perform the force measurement using a JPK NanoWizard® 4 NanoScience AFM and JPK NanoWizard® Control Software v6.1.115 coupled to an optical microscope that allows observation of the sample. Force measurements were subsequently performed first in the quantitative imaging mode, using the following parameters: a contact setpoint of 0.5 nN, a z length of 1 μm, a constant approach and retraction speed of 50 μm/s and a scan area of 50 μm × 50 μm (256 × 256 pixels) on top of a corneocyte. A conventional force volume map was also recorded on a smaller area (500 nm × 500 nm, 16 × 16 pixels) with a contact setpoint of 0.25 nN and a slower constant approach and retraction speed of 1 μm/s. Data were analyzed using the data analysis software from JPK and Origin software.

## Acknowledgements

The authors gratefully acknowledge expert advice and support from Drs. Frank Murphy, Jesse Hopkins, and Srinivas Chakravarthy from Argonne National Laboratory. Y.F.D. thanks Dr. Joan Geoghegan for providing corneocytes for AFM analysis. This research was conducted with NIH funding through NIGMS (R01 GM094363 to ABH) and NIAID (R01 AI162964 to PDF, ARH, and ABH; R01 AI153185 to ARH; and R01 AI139479 to NCF). Work at UCLouvain was supported by the European Research Council (ERC) under the European Union’s Horizon 2020 research and innovation programme (grant agreement n°693630), and the National Fund for Scientific Research (FNRS). Glycan array screening was carried out by the Protein-Glycan Interaction Resource of the CFG (supported by NIGMS grant R24 GM098791) and the National Center for Functional Glycomics (NCFG) at Beth Israel Deaconess Medical Center, Harvard Medical School (supported by NIGMS grant P41 GM103694). This research used resources of the Advanced Photon Source, a U.S. Department of Energy (DOE) Office of Science User Facility operated for the DOE Office of Science by Argonne National Laboratory under Contract No. DE-AC02-06CH11357. Diffraction data were collected at the Northeastern Collaborative Access Team (NE-CAT) beamlines, which are funded by the NIH through NIGMS (P30 GM124165). The Eiger 16M detector on 24-ID-E is funded by a NIH-ORIP HEI grant (S10OD021527). SEC-MALS-SAXS data were collected at the Biophysics Collaborative Access Team (BioCAT) beamline, supported by NIH grant P41 GM103622 through NIGMS. Use of the Pilatus 3 1M detector was provided by grant 1S10OD018090 from NIGMS. Brpt1.5 SAXS data collection was conducted at the Advanced Light Source (ALS), a national user facility operated by Lawrence Berkeley National Laboratory on behalf of the Department of Energy, Office of Basic Energy Sciences, through the Integrated Diffraction Analysis Technologies (IDAT) program, supported by DOE Office of Biological and Environmental Research. Additional support comes from the NIH project ALS-ENABLE (P30 GM124169) and a High-End Instrumentation Grant S10OD018483. The content is solely the responsibility of the authors and does not necessarily reflect the official views of the National Institute of General Medical Sciences or the National Institutes of Health.

## Author Contributions

JJM, CC, KBM, AEY, CTC, NCF, MMG, YFD, PDF, ARH, and ABH designed experiments; JJM, CC, KBM, RY, AEY, CTC, and DV collected data; all authors analyzed data; and JJM, CC, KBM, RY, AEY, NCF, MMG, YFD, PDF, ARH, and ABH wrote the article.

