## Supplementary Information for "Mechanistic basis of staphylococcal interspecies competition for skin colonization"

Figures:
Figure S1. Comparison of Aap and SasG: domain architecture and lectin sequence.
Figure S2. Analytical ultracentrifugation of Aap and SasG.
Figure S3. Structural models of Aap-Brpt from SEC-SAXS.
Figure S4. R1 and R2 relaxation data on Aap lectin.
Figure S5. Chemical shift perturbation data for lactosamine titration of Aap lectin.
Figure S6. Aap mediates adhesion to human corneocytes.
Figure S7. Aap mutants do not engage human corneocytes.

Tables:
Table S1. Crystallographic data collection and refinement statistics.
Table S2. Strains used in study.

a

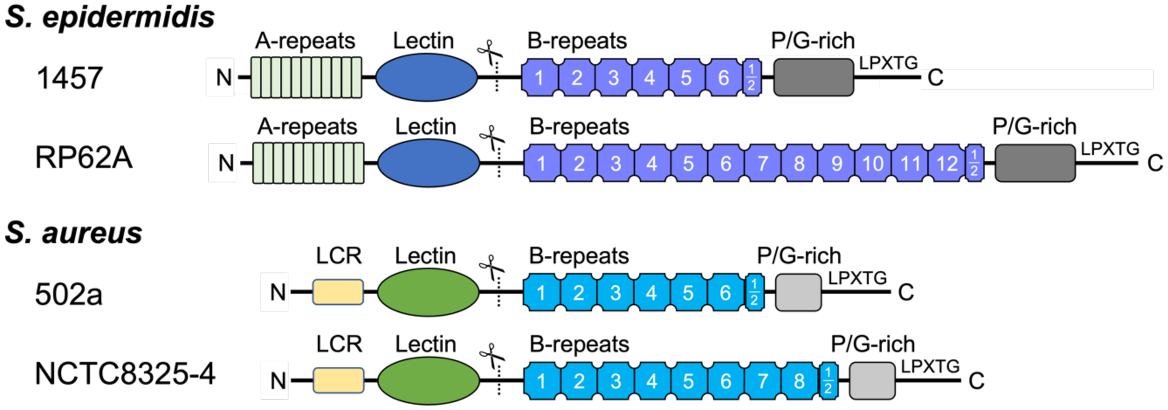


b

```
Aap     IPPTTVKGRDNYDFYGRVDIESNPTDLNATNLTRYNYGQPPGTTTAGAVQFKNQVSFDKDFDFNIRVANN
SasG    T-PITIQGKEHFEGYGSVDIQKNPTDLGVSEVTRFNVGNESN-GLIGALQLKNKIDFSKDFNFKVRVANN
          * *::*::::: ** ***:.*****..:::**:* *: ..     **:*:**::.*.***:*::*****

Aap     RQSNTTGADGWGFMFSKKDGDDFLKNGGILREKGTPSAAGFRIDTGYYNNDPLDKIQKQAGQGYRGYGTF
SasG    HQSNTTGADGWGFLFSKGNAEEYLTNGGILGDKGLVNSGGFKIDTGYIYTSSMDKTEKQAGQGYRGYGAF
        :************:*** :.:::*.***** :**  .:.**:*****  ...:** :***********:*

Aap     VKNDSQGNTSKVGSGT--PSTDFLNYADNTTNDLDGKFHGQKLNNVNLKYNASNQTFTATYAGKTWTATL
SasG    VKNDSSGNSQMVGENIDKSKTNFLNYADNSTNTSDGKFHGQRLNDVILTYVASTGKMRAEYAGKTWETSI
        *****.**:. **..   ..*:*******:**  *******:**:* *.* **. .: * ****** :::

Aap     SELGLSPTDSYNFLVTSSQYGNGNSGTYASGVMRADLDGATLTYTPK-
SasG    TDLGLSKNQAYNFLITSSQRWGLNQGINANGWMRTDLKGSEFTFTPEA
        ::**** .::****:****  . *.*  *.* **:**.*::*:**:
```

**Fig. S1: Comparison of Aap and SasG: domain architecture and lectin sequence.** (**a**) Domain architecture of Aap and SasG variants from different strains of *S. epidermidis* and *S. aureus*, respectively. (**b**) Sequence alignment of Aap and SasG lectin domains.

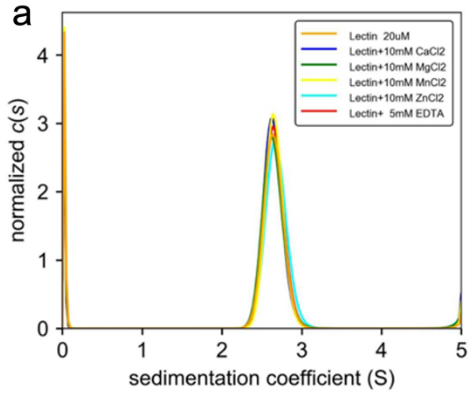


| Sample | s (Svedbergs) | $f/f_0$ | Molar mass (kDa) |
|---|---|---|---|
| **Aap lectin** | 2.66 | 1.30 | 29.6 |
| **Lectin +$Ca^{2+}$** | 2.65 | 1.32 | 30.4 |
| **Lectin +$Mg^{2+}$** | 2.65 | 1.32 | 30.2 |
| **Lectin +$Mn^{2+}$** | 2.67 | 1.28 | 29.2 |
| **Lectin +$Zn^{2+}$** | 2.69 | 1.31 | 30.8 |
| **Lectin +EDTA** | 2.67 | 1.30 | 29.9 |

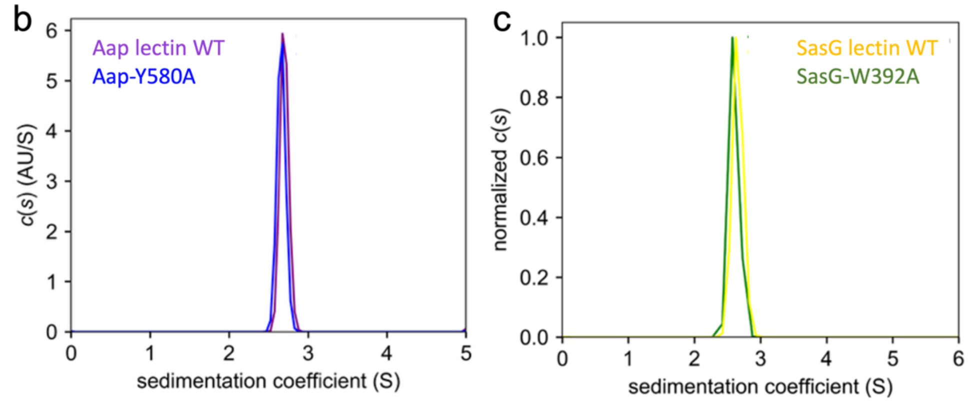


**Fig. S2: Analytical ultracentrifugation of Aap and SasG.** (a) Sedimentation velocity analytical ultracentrifugation reveals that Aap-lectin sediments as a monomer even in the presence of various divalent metal ions. (b) Comparison of the sedimentation coefficient distributions for WT Aap-lectin versus the Aap-Y580A lectin mutant, confirming that the mutant domain has a similar fold and assembly state. (c) Comparison of the sedimentation coefficient distributions for WT SasG-lectin versus the SasG-W392A lectin mutant, confirming that the mutant domain has a similar fold and assembly state.

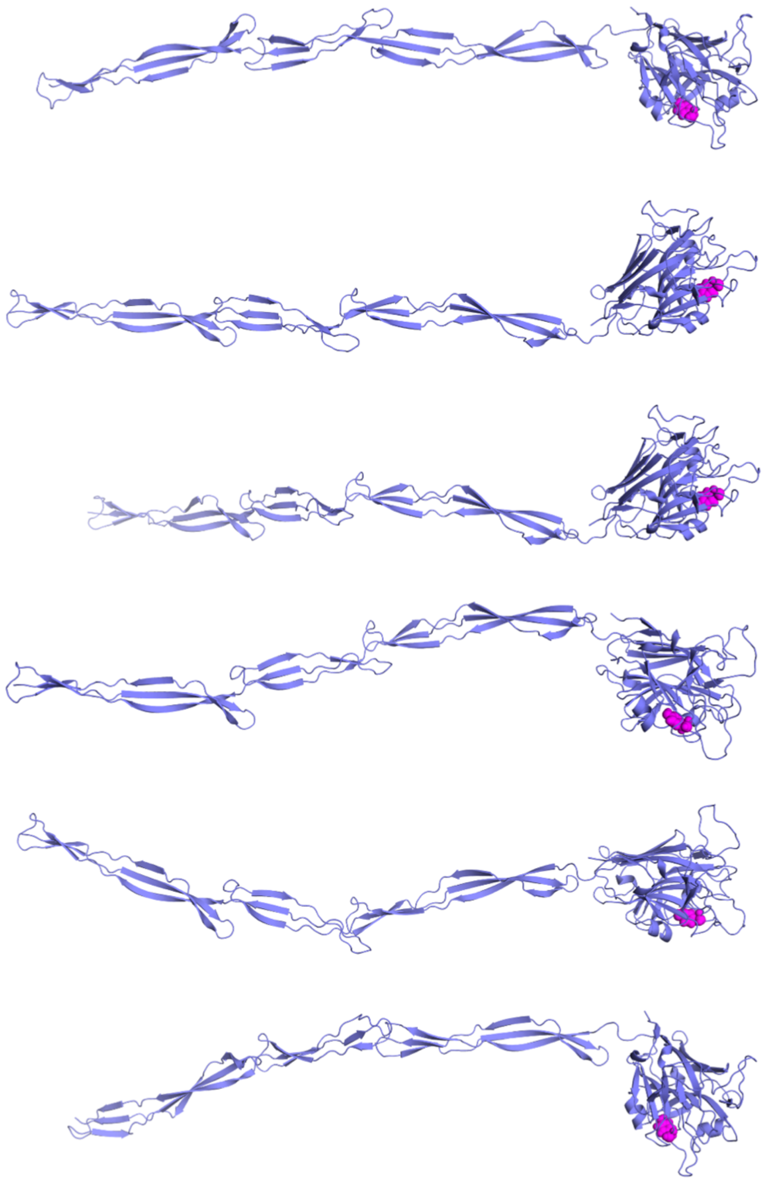

**Fig. S3: Structural models of Aap-Brpt from SEC-SAXS.** Comparison of the best-fit Aap-Brpt models (from a pool generated by SASSIE) determined by CRYSOL to be most consistent with the experimental scattering curve.

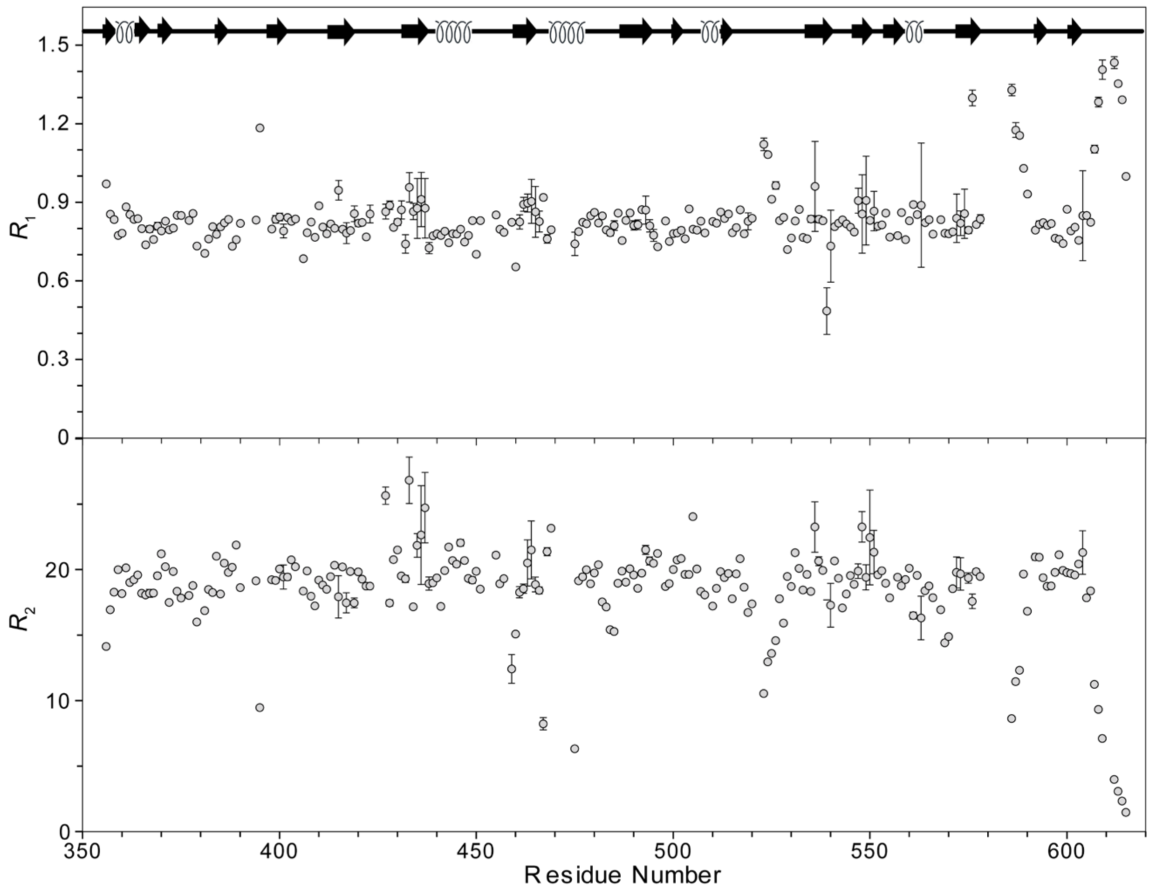


**Fig. S4: $R_1$ and $R_2$ relaxation data on Aap lectin.** $R_1$ (top); $R_2$ (bottom) values of Aap-lectin determined at 298 K in the 600 MHz Bruker spectrometer.

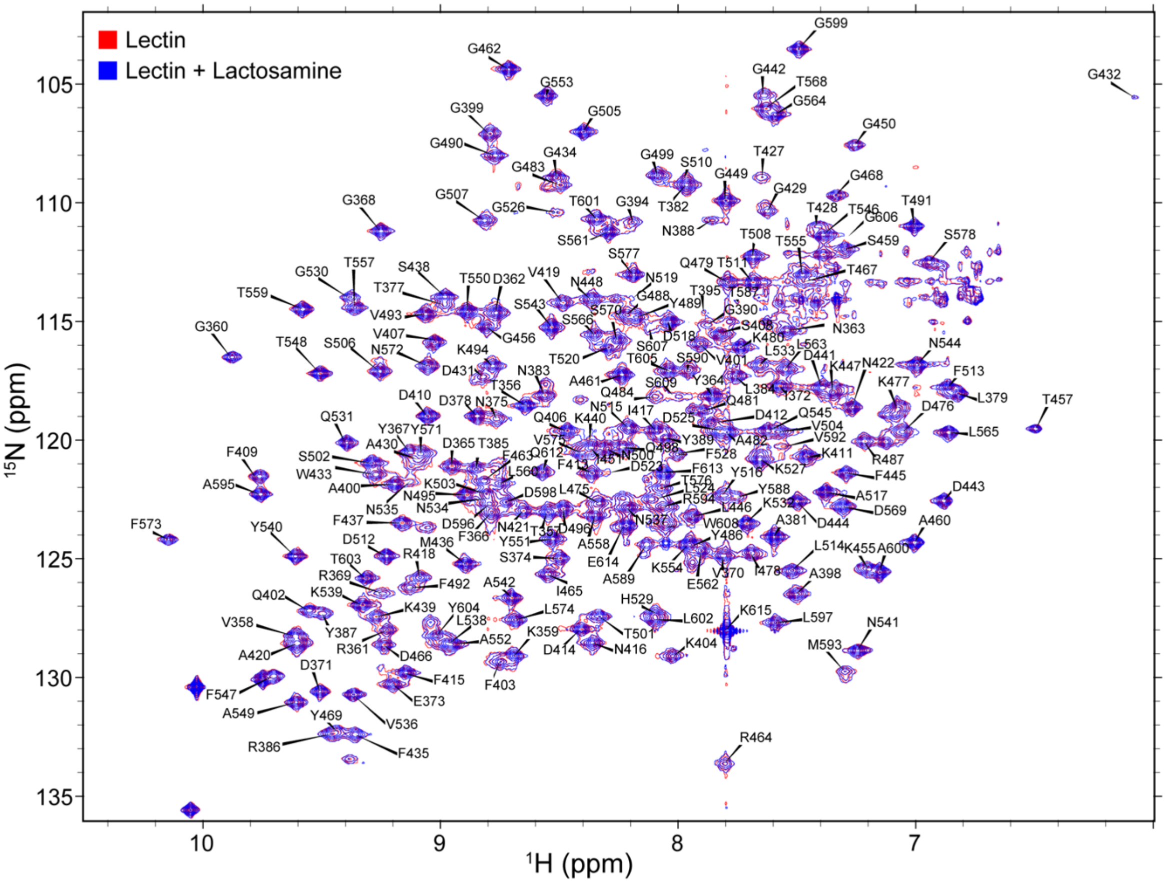


**Fig. S5: Chemical shift perturbation data for lactosamine titration of Aap lectin.** Overlay of $^{15}$N TROSY-HSQC of 0.2 mM Aap-lectin (red) and 0.2 mM Aap-lectin + 20 mM lactosamine (blue). No significant chemical shift perturbation was observed upon the addition of lactosamine, which could be due to the low affinity of lactosamine for the Aap-lectin.

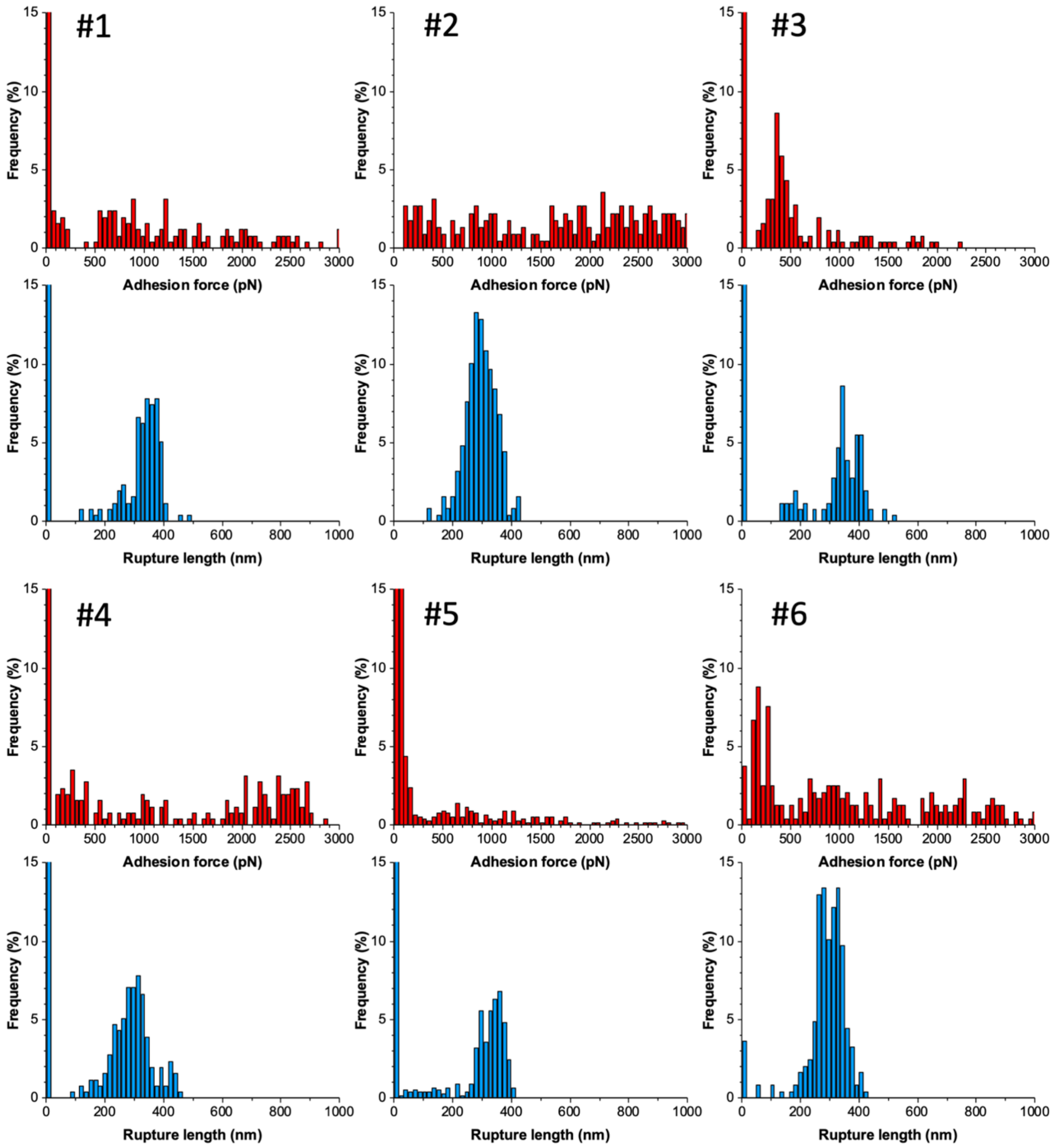


**Fig. S6: Aap mediates adhesion to human corneocytes.** Adhesion force (red) and rupture length (blue) histograms obtained by recording force-distance curves between 6 different *S. epidermidis* 1457 *ΔsepAΔica* - human corneocyte pairs (n= 256 curves for each pair).

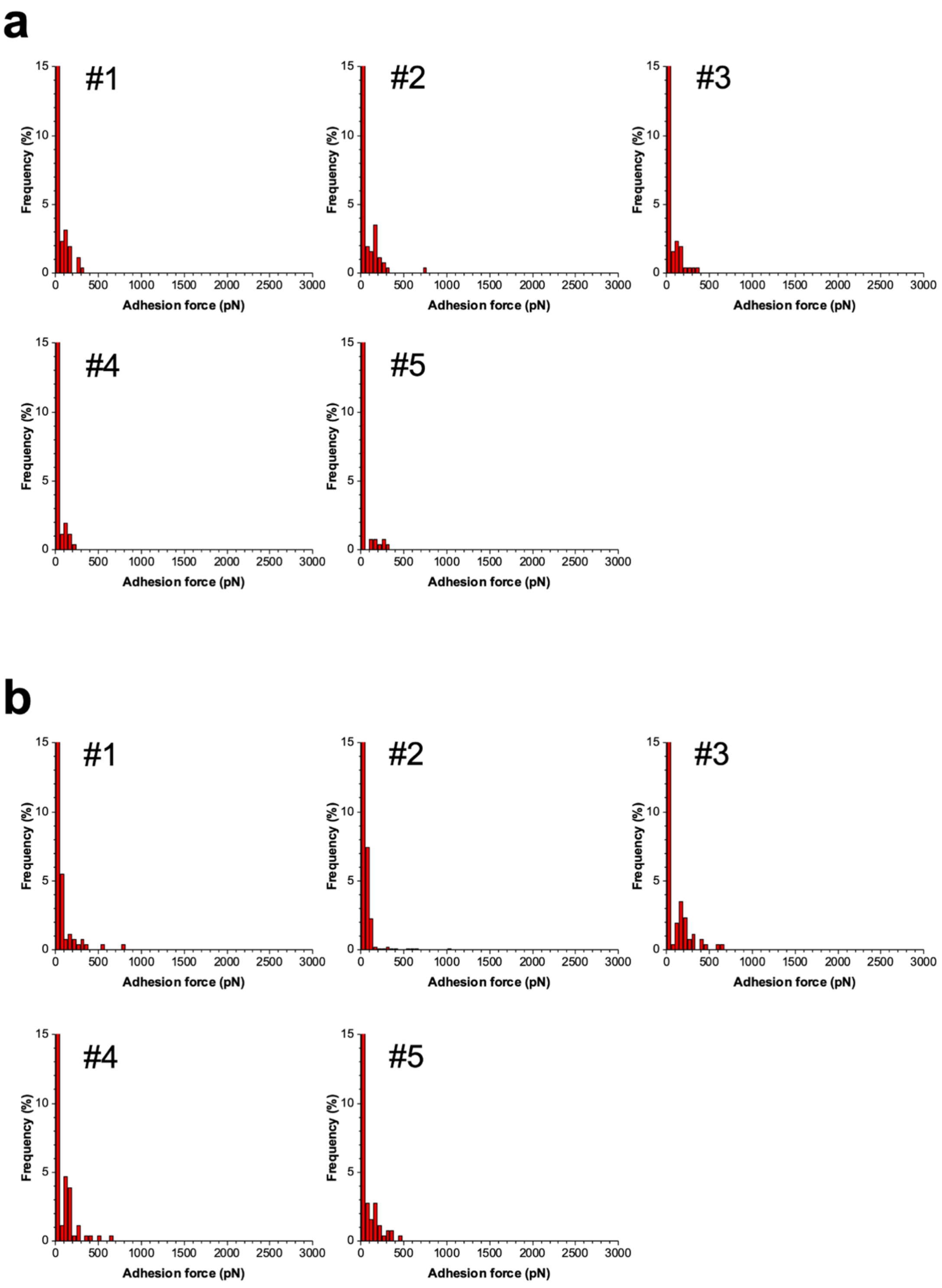


**Fig. S7: Aap mutants do not engage human corneocytes.** Adhesion force (red) and rupture length (blue) histograms obtained by recording force-distance curves between (**a**) 5 different *S. epidermidis* 1457 *ΔaapΔica* - human corneocyte pairs (n= 256 curves for each pair) and (**b**) 5 different *S. epidermidis* 1457 *ΔsarAΔica* - human corneocyte pairs (n= 256 curves for each pair).

**Table 1. Crystallographic data collection and refinement statistics.**

| | **Aap-lectin (*S. epidermidis*)** | **SasG-lectin (*S. aureus*)** |
|---|---|---|
| Wavelength (Å) | 0.97780 | 0.97872 |
| Resolution range (Å)[a] | 34.29–1.06 (1.10–1.06) | 33.63–1.93 (2.0–1.93) |
| Space group | $P2_12_12_1$ | C2 |
| Unit cell dimensions | | |
| *a, b, c* (Å)<br>*α, β, γ* (°) | 35.66, 68.57, 109.27<br>90, 90, 90 | 127.10, 43.50, 39.40<br>90, 90.56, 90 |
| Total reflections (unique reflections) | 1,005,349 (88,819) | 187,090 (18,690) |
| Unique reflections | 121,879 (11,899) | 16,284 (1,613) |
| Multiplicity | 8.2 (7.5) | 11.5 (11.6) |
| Completeness (%) | 99.65 (98.48) | 99.03 (98.59) |
| Mean I/sigma(I) | 14.88 (4.39) | 141.31 (25.00) |
| Wilson B-factor | 8.33 | 11.20 |
| R-merge | 0.091 (0.554) | 0.591 (0.689) |
| R-meas | 0.097 (0.596) | 0.621 (0.723) |
| R-pim | 0.034 (0.216) | 0.191 (0.218) |
| CC1/2 | 0.997 (0.911) | 0.801 (0.756) |
| CC* | 0.999 (0.976) | 0.943 (0.928) |
| Reflections used in refinement | 121,841 (11,890) | 16,263 (1,613) |
| Reflections used for R-free | 2,000 (196) | 1,629 (166) |
| R-work | 0.117 (0.131) | 0.148 (0.157) |
| R-free | 0.126 (0.147) | 0.199 (0.237) |
| CC (work) | 0.976 (0.965) | 0.914 (0.854) |
| CC (free) | 0.971 (0.978) | 0.908 (0.740) |
| Number of non-hydrogen atoms | 2380 | 2098 |
| macromolecules | 2010 | 1921 |
| ligands | 14 | 1 |
| solvent | 356 | 176 |
| protein | 252 | 255 |
| RMS (bonds) | 0.006 | 0.007 |
| RMS (angles) | 1.02 | 0.79 |
| Average B-factor | 11.69 | 12.16 |
| macromolecules | 9.75 | 11.59 |
| ligands | 23.56 | 4.94 |
| solvent | 22.17 | 18.49 |
| Ramachandran favored (%) | 97.60 | 99.21 |
| Ramachandran allowed (%) | 2.40 | 0.79 |
| Ramachandran outliers (%) | 0.00 | 0.00 |
| Rotamer outliers (%) | 1.86 | 0.00 |
| MolProbity clashscore (percentile) | 2.81 ($93^{rd}$) | 1.63 ($100^{th}$) |

[a] Values in parentheses are for the highest resolution shell.

**Table S2. Strains used in study**

| Bacterial strain | Relevant genotype/phenotype | Source |
|---|---|---|
| *S. epidermidis* 1457 Δ*ica*/pCM29 | 1457*icaADBC::dhfr*; PIA-negative, Aap-positive; $Tmp^R$. Plasmid pCM29 ($Chl^R$) encodes sGFP driven by the constitutive promoter *sarA* P1. | (1-3) |
| *S. epidermidis* 1457 Δ*ica* Δ*aap* /pCM29 | 1457*icaADBC::dhfr aap::tetM*; PIA-negative, Aap-negative; $Tmp^R$, $Tet^R$. Carries pCM29 ($GFP^+$; $Chl^R$) | (1-3) |
| *S. epidermidis* 1457 Δ*ica* Δ*sepA*/pCM29 | 1457*icaADBC::dhfr sepA* PIA-negative, full length Aap (A-domain not cleaved) due to lack of processing by SepA; $Tmp^R$. Carries pCM29 ($GFP^+$; $Chl^R$) | (4) |
| *S. epidermidis* 1457 Δ*ica* Δ*sarA*/pCM29 | 1457*icaADBC::dhfr sarA::tetM*; PIA-negative, Aap A-domain negative due to unregulated SepA production due to *sarA* mutation; $Tmp^R$, $Tet^R$. Carries pCM29 ($GFP^+$; $Chl^R$) | (4) |
| *S. carnosus* TM300 pALC2073 | *S. carnosus* TM300 ΔSCA_2092 ΔSCA_2238::sGFP. Carries pALC2073 ($Chl^R$) | (5) |
| *S. carnosus* TM300 pALC2073-$SasG_{COL}$ | *S. carnosus* TM300 ΔSCA_2092 ΔSCA_2238::sGFP. Carries pALC2073-$SasG_{COL}$ ($Chl^R$) | (5) |

Abbreviations: Tmp (Trimethoprim); Chl (Chloramphenicol); Tet (Tetracycline).
